# Parkinsonism Sac domain mutation in Synaptojanin-1 affects ciliary properties in iPSC-derived dopaminergic neurons

**DOI:** 10.1101/2023.10.12.562142

**Authors:** Nisha Mohd Rafiq, Kenshiro Fujise, Martin Shaun Rosenfeld, Peng Xu, Yumei Wu, Pietro De Camilli

**Author notes:** Interfaculty Institute of Biochemistry, DZNE Tübingen, University of Tübingen, Auf der Morgenstelle 34, 72076 Tübingen, Germany. Address correspondence to: Nisha Mohd Rafiq; Pietro De Camilli.

## Abstract

Synaptojanin-1 (SJ1) is a major neuronal-enriched PI(4,5)P^2^ 4- and 5-phosphatase implicated in the shedding of endocytic factors during endocytosis. A mutation (R258Q) that impairs selectively its 4-phosphatase activity causes Parkinsonism in humans and neurological defects in mice (SJ1^RQ^KI mice). Studies of these mice showed, besides an abnormal assembly state of endocytic factors at synapses, the presence of dystrophic nerve terminals selectively in a subset of nigro-striatal dopamine (DA)-ergic axons, suggesting a special lability of DA neurons to the impairment of SJ1 function. Here we have further investigated the impact of SJ1 on DA neurons using iPSC-derived SJ1 KO and SJ1^RQ^KI DA neurons and their isogenic controls. In addition to the expected enhanced clustering of endocytic factors in nerve terminals, we observed in both SJ1 mutant neuronal lines increased cilia length. Further analysis of cilia of SJ1^RQ^DA neurons revealed abnormal accumulation of the Ca^2+^ channel Ca_v_1.3 and of ubiquitin chains, suggesting an impaired clearing of proteins from cilia which may result from an endocytic defect at the ciliary base, where a focal concentration of SJ1 was observed. We suggest that SJ1 may contribute to the control of ciliary protein dynamics in DA neurons, with implications on cilia-mediated signaling.

## INTRODUCTION

While the cause of most Parkinson’s disease (PD) is not known, mutations in a selected list of genes are responsible for the development of familial forms of the disease, often Early-Onset Parkinsonism (EOP)(Blauwendraat et al. 2020). One such gene is *SYNJ1*, which encodes the protein synaptojanin-1 (SJ1), a polyphosphoinositide phosphatase highly expressed in neurons and enriched at synapses(McPherson et al. 1996, Krebs et al. 2013, Quadri et al. 2013). SJ1 dephophorylates PI(4,5)P_2_ via the sequential action of two tandemly arranged inositol phosphatase modules: a central 5-phosphatase domain and an N-terminal Sac1 domain which functions primarily as a 4-phosphatase (McPherson et al. 1996, Guo et al. 1999, Nemoto et al. 2000). These catalytic modules are followed by a proline-rich region which is responsible for its subcellular targeting and undergoes alternative splicing to generate a shorter (145 kD, the predominant neuronal form) and a longer (170 kD) isoform (McPherson et al. 1996, Ramjaun and McPherson 1996, Rosivatz 2006). One of the main known roles of SJ1 is to participate in the shedding from endocytic vesicles of clathrin coats and other endocytic factors, including actin regulatory proteins, which bind PI(4,5)P_2_ at the plasma membrane to initiate the endocytic reaction(Cremona et al. 1999, Di Paolo and De Camilli 2006). While absence of SJ1 leads to early postnatal lethality in mice (Cremona et al. 1999) and humans (Dyment et al. 2015, Hardies et al. 2016), a patient R258Q missense mutation (SJ1^RQ^) (accession number: NM_003895) also known as R219Q (accession number: NM_001160302) is responsible for EOP with epilepsy. This mutation selectively abolishes the catalytic action of its Sac1 domain (SJ1^RQ^)(Krebs et al. 2013). We previously showed that knock-in mice with this mutation (SJ1^RQ^KI) display neurologic manifestations reminiscent of those of human patients (Cao et al. 2017). These manifestations are accompanied at the cellular level not only by endocytic defects and an accumulation of clathrin-coated vesicles at synapses, but also by degenerative changes selectively of a subset of dopaminergic nerve terminals in the dorsal striatum (Cao et al. 2017, Ng et al. 2023).

One cell compartment which is regulated by PI4P and PI(4,5)P_2_ dynamics is the primary cilium (Chavez et al. 2015, Garcia-Gonzalo et al. 2015, Demmel et al. 2016, Ojeda Naharros and Nachury 2022). PI(4,5)P_2_ in the plasma membrane of the ciliary pocket at the base of the cilium, which is a site of intense exo-endocytosis, helps regulate the turnover of cilia-related signaling proteins (Garcia-Gonzalo et al. 2015, Demmel et al. 2016, Nachury and Mick 2019). Moreover PI(4,5)P_2_ is the precursor of the pool of PI4P generated in the ciliary shaft through dephosphorylation of PI(4,5)P_2_ by INPP5E, a polyphosphoinositide 5-phosphatase concentrated in the shaft of primary cilia. This PI4P pool has a critical role in cilia biology (Bielas et al. 2009, Jacoby et al. 2009, Chavez et al. 2015, Garcia-Gonzalo et al. 2015, Phua et al. 2017, Klink et al. 2020). Primary cilia are key players in the hedgehog signaling pathway which has a crucial importance in the nigrostriatal system (Hynes et al. 1995, Nordstroma et al. 2015, Dhekne et al. 2018, Derderian et al. 2023). The importance of hedgehog signaling in the development of DA neurons is proven by the essential requirement of Sonic Hedgehog (Shh) for the differentiation of iPSCs into DA neurons (Kriks et al. 2011, Kim et al. 2021). Primary cilia of neurons are increasingly recognized as major signaling hub with a major impact on neuronal function. Interestingly, disease causing mutations in another PD gene, LRRK2 (PARK8)(Paisan-Ruiz et al. 2004, Zimprich et al. 2004, Poewe et al. 2017, Alessi and Sammler 2018, Sobu et al. 2021) interfere with ciliogenesis (Dhekne et al. 2018, Khan et al. 2021, Sobu et al. 2021), suggesting a potential contribution of ciliary-related defects to PD pathology. While one effect of PD LRRK2 mutations is to impact DA neurons indirectly, via an impairment of cilia-dependent hedgehog signaling in striatal cholinergic neurons (Dhekne et al. 2018, Khan et al. 2021), additional directs effects of these mutations via an impairment of cilia in DA neurons cannot be excluded. These considerations raise the question of whether phenotypic manifestations of SJ1 impairment may include perturbations of ciliary functions and whether such perturbations may occur in DA neurons.

Here we have used iPSC-derived DA neurons as a model system to address this question. We report that DA neurons with impaired SJ1 function have abnormally long cilia which display an ectopic accumulation of ubiquitinated proteins within them. The Ca_v_1.3, a voltage-gated calcium channel, which is important for the rhythmic pacemaking activity of DA neurons (Gregory et al. 2011, Felix and Weiss 2017, Liss and Striessnig 2019, Grimaldo et al. 2022), is also ectopically accumulated within them. Together, our results demonstrate a role of SJ1 in the cilia of DA neurons and implicates this protein in the control of their signaling properties.

## RESULTS

### Generation of WT and SJ1 mutant iPSC-derived DA neurons

Human iPSCs (WTC11 line) were gene edited in house by CRISPR/Cas9 to delete expression of SJ1 (SJ1 KO). Correct editing was validated by PCR and absence of SJ1 in KO cells was confirmed by western blotting (Supplementary Figure 1A and B). iPSCs (KOLF2.1 line) harboring the EOP RQ mutation at position 258 (accession number: NM_003895) were obtained from the iPSC Neurodegeneration Initiative (iNDI)(Ramos et al. 2021) and validated by polymerase chain reaction (PCR). SJ1 KO and SJ1^RQ^KI iPSCs, as well as their corresponding isogenic controls were differentiated either into cortical-like i^3^neurons or into DA neurons (Supplementary Figure 1C-G). To generate cortical-like i^3^Neurons, we used the doxycycline-inducible neurogenin-2 (NGN2)-driven differentiation protocol(Wang et al. 2017) as described in Fernandopulle et al. (2018) which results in mature neuronal cultures within 15-19 days. For the generation of DA neurons, we used the procedure described by Kriks et al. (2011) and Bressan et al. (2021). This differentiation process is slower than the NGN2-driven neuronal differentiation (Fernandopulle et al. 2018, Bressan et al. 2021, Hulme et al. 2022). However, 30 days from the beginning of differentiation, cells had acquired neuronal morphology with the formation of a complex network of processes (Supplementary Figure 1C and D). Moreover, western blotting and immunofluorescence of these cultures showed the expression of two key markers of DA neurons, tyrosine hydroxylase (TH) and the dopamine transporter (DAT), in both the control and the two SJ1 mutant lines (Supplementary Figure 1E-I).

### iPSC-derived SJ1 mutant DA neurons display abnormal accumulation of endocytic factors in nerve terminals

A key and defining phenotype of SJ1 KO and SJ1^RQ^KI neurons *in situ* and in primary cultures is a very robust and exaggerated accumulation in their nerve terminals of endocytic membrane intermediates and endocytic factors, including clathrin coat components and their accessory factors, with amphiphysin-2 being the most strikingly accumulated protein (Cao et al. 2017). To validate the use of iPSC-derived DA neurons as model systems to assess the impact of SJ1 mutations, we examined if this phenotype was recapitulated in these cells.

At day 50-55 from the beginning of differentiation, SJ1 KO neurons, SJ1^RQ^KI DA neurons and their corresponding control neurons showed a similar and prominent punctate pattern of immunoreactivity for the synaptic vesicle marker synaptophysin, revealing abundant formation of synapses in all four conditions. However, a very strong and robust accumulation of puncta of amphiphysin-2 immunoreactivity, which overlapped with synaptophysin immunoreactivity (Figure 1A-F), was observed in SJ1 KO and SJ1^RQ^KI DA neurons, but not in control neurons, demonstrating that the accumulation of endocytic factors typical of SJ1 KO neurons is replicated in these iPSC-derived neurons. These accumulations were also seen when SJ1^RQ^KI DA neurons were cocultured for 7 days with iPSC-derived medium spiny neurons (MSNs) (Figure 1G-J) using a microfluidic compartmentalization device (eNuvio). In this device, DA neurons and MSNs are seeded in two distinct chambers connected by narrow channels through which axons can grow. Large abnormal puncta of amphiphysin-2 immunoreactivity, which overlapped with puncta positive for synapsin, a marker of presynaptic nerve terminals (De Camilli et al. 1983), were observed in both chambers, with the puncta found in the MSN-containing chamber likely reflecting DA synapses on MSNs. We conclude that iPSC-derived DA neurons are good models to study SJ1 mutant phenotypes.

**Figure 1:**
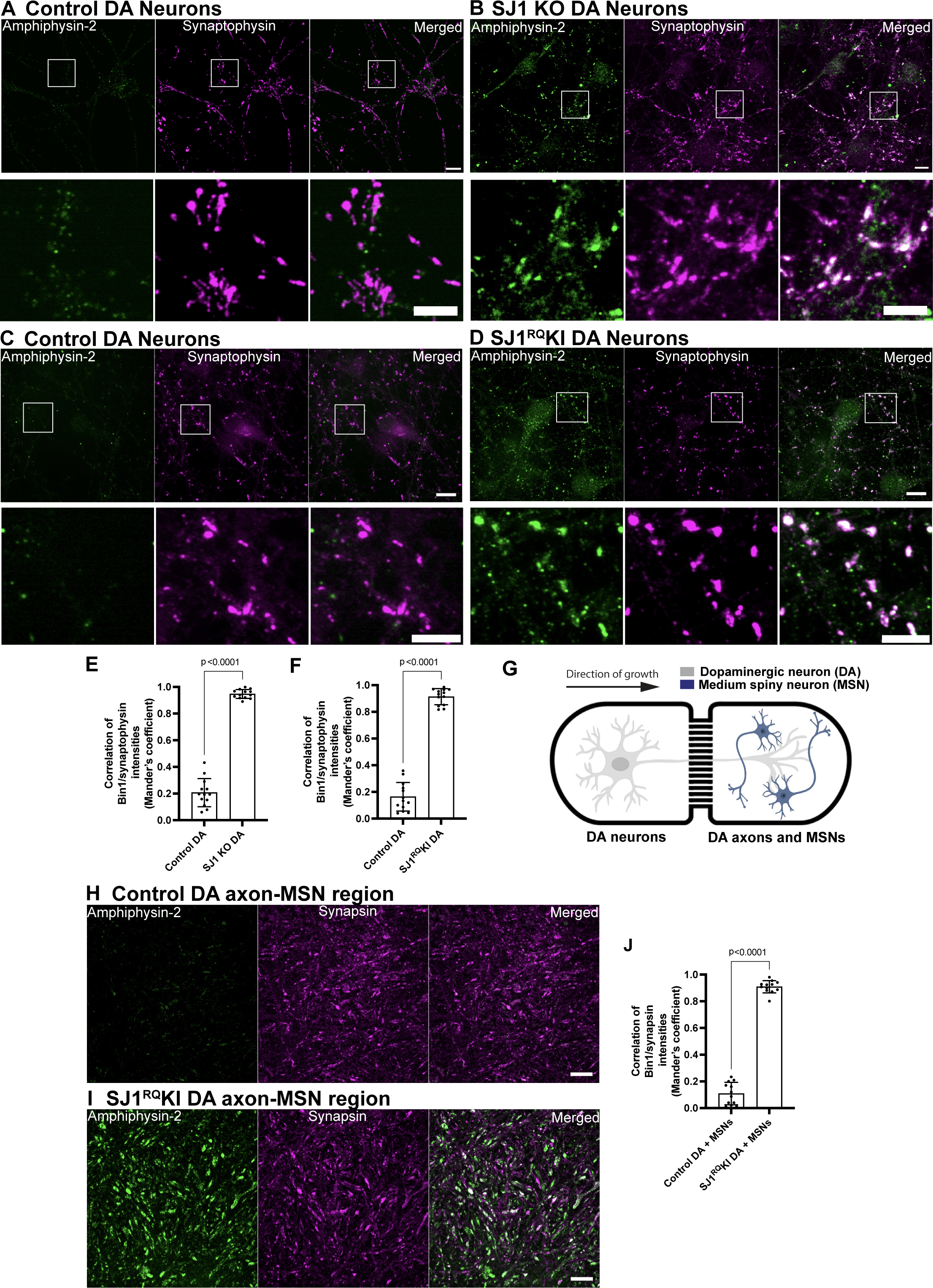
SJ1 KO and SJ1^RQ^KI iPSC-derived DA neurons show presynaptic clustering of amphiphysin-2. (A-D) Fluorescence images of control (A, C), SJ1 KO (B) and SJ1^RQ^KI (D) DA neurons (day 50-55) immunolabeled with antibodies directed against amphiphysin-2 (green) and synaptophysin, a presynaptic marker, (magenta). SJ1 KO neurons and the corresponding controls are derived WTC11 iPSCs, while SJ1^RQ^KI neurons and corresponding controls are derived from KOLF2.1 iPSCs (Scale bars, 10 µm). High magnifications of boxed areas are shown below each panel (Scale bars, 5 µm). Note the striking enhancement of amphiphysin-2 immunoreactivity that overlaps with synaptophysin-positive structures in SJ1 KO and SJ1^RQ^KI DA neurons, relative to controls. (E and F) Quantification of amphiphysin-2 clustering intensities shown in A-D, represented as mean ± S.D., pooled from at least two independent experiments (n ≥ 10 cells per experiment). (G) Diagram showing a schematic view of iPSC-derived DA (day 55) and iPSC-derived medium spiny neurons (MSNs) (from Brainxell cells, day 7 post-thaw) co-cultured in the microfluidic device. (H and I) Immunofluorescence images of amphiphysin-2 (green) and synapsin (magenta) immunoreactivities in the MSN containing chamber of neuronal co-cultures generated with control (H) or SJ1^RQ^KI DA neurons (I) (Scale bars, 10 µm). (J) Quantification of fluorescence intensity of amphiphysin-2 puncta in the MSN containing chamber (mean ± S.D. from two independent experiments; n ≥ 20 regions per experiment).

### Presence of primary cilia in iPSC-derived DA neurons and abnormal ciliary length in SJ1 KO and SJ1^RQ^KI DA neurons

Cilia brightly positive for the primary cilia marker Arl13b (Caspary et al. 2007) were clearly visible in undifferentiated iPSCs, but no longer detectable after differentiation to cortical-like i^3^Neurons (Figure 2A and Figure 2B). This is in agreement with the decrease of the levels of mRNAs encoding cilia-related proteins as detected by RNAseq during iPSC-differentiation in i^3^Neurons (Tian et al. 2019). In contrast, the great majority of iPSC-derived DA neurons retained Arl13b -positive cilia (89.45 ± 1.68%; mean ± S.E.M.), which were also positive for acetylated tubulin (a general cilia marker) and for adenylate cyclase type III (AC3), a marker specific of neuronal cilia (Sipos et al. 2018, Sterpka and Chen 2018)(Figure 2C-E).

**Figure 2:**
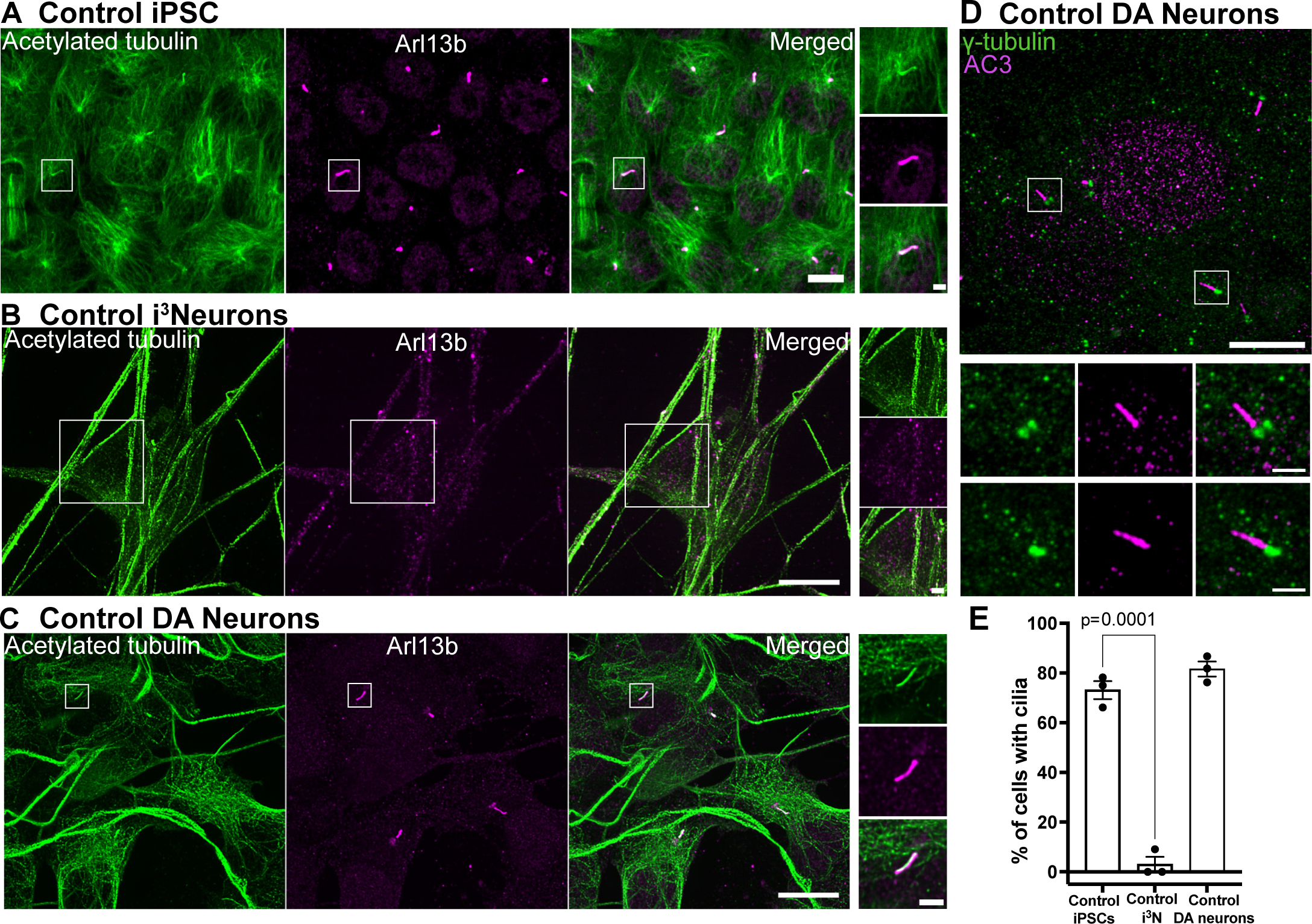
iPSC-derived DA neurons have primary cilia. (A-C) Fluorescence images of undifferentiated iPSCs (A), i^3^Neurons (day 19, B) and iPSC-derived DA neurons (day 30, D)(all from KOLF2.1 iPSCs) immunolabeled with antibodies directed against acetylated α-tubulin (green) and Arl13b (magenta)(Scale bars, 10 μm). High magnification images of the boxed areas in A-C are shown on the right (Scale bars, 2 µm). iPSCs have primary cilia but cilia are no longer present in i^3^Neurons, while they are still present in DA neurons. (D) Fluorescence images of DA neurons immunolabeled with antibodies against γ-tubulin (green) and the neuronal-specific primary cilia marker, adenylate cyclase type III (AC3, magenta), confirming the neuronal properties of these neurons. (F) Percentage of cells with cilia (mean ± S.E.M.) from three independent experiments; n ≥ 20 cells per experiment).

Cilia, as assessed by Arl13b, acetylated microtubules and AC3 immunolabeling, were almost two-fold longer in SJ1 KO neurons when compared to control neurons, while the percentage of cilia-forming cells was the same in both conditions (Figure 3A-F). Furthermore, abnormally shaped Arl13b-positive cilia were observed in SJ1 KO DA neurons with presence of multiple abnormal cilia in a small proportion of SJ1 KO DA neurons, but not in their controls (Supplementary Figure 2).

**Figure 3:**
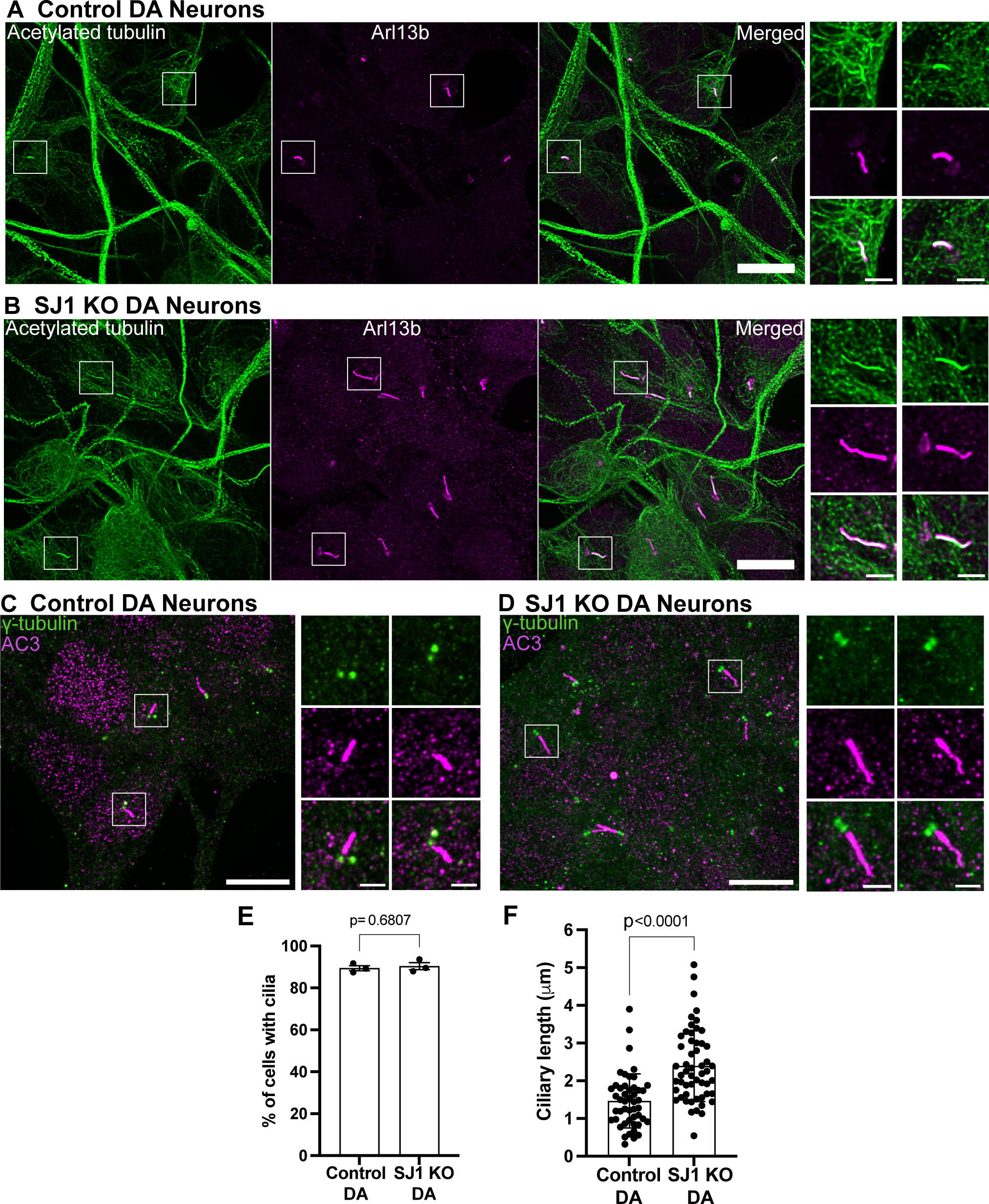
Abnormal ciliary length in SJ1 KO iPSC-derived DA neurons relative to control iPSC-derived DA neurons. (A-D) Fluorescence images of control (A and B) and SJ1 KO (C and D) DA neurons (day 30) immunolabeled with antibodies against acetylated α-tubulin (green), Arl13b (magenta) or γ-tubulin (green) and the neuronal-specific primary cilia marker, AC3 (magenta)(Scale bars, 10 µm). High magnification of the boxed areas in A-D are shown on the right of each panel (Scale bars, 2 µm). (E-F) Percentage of ciliated cells (E) and cilia length (F) of control and SJ1 KO DA neurons represented as mean ± S.D. (data pooled from three independent experiments; n ≥ 10 cells per experiment).

We next analyzed presence of cilia in two different iPSC-derived clones of SJ1^RQ^KI DA neurons (Figure 4). While again there was no difference in the percentage of cilia-forming DA neurons relative to controIs, the length of cilia was significantly longer in both clones in comparison to control (Figure 4A-E). We conclude that lack of a functional SJ1 affects some properties of cilia in DA neurons.

**Figure 4:**
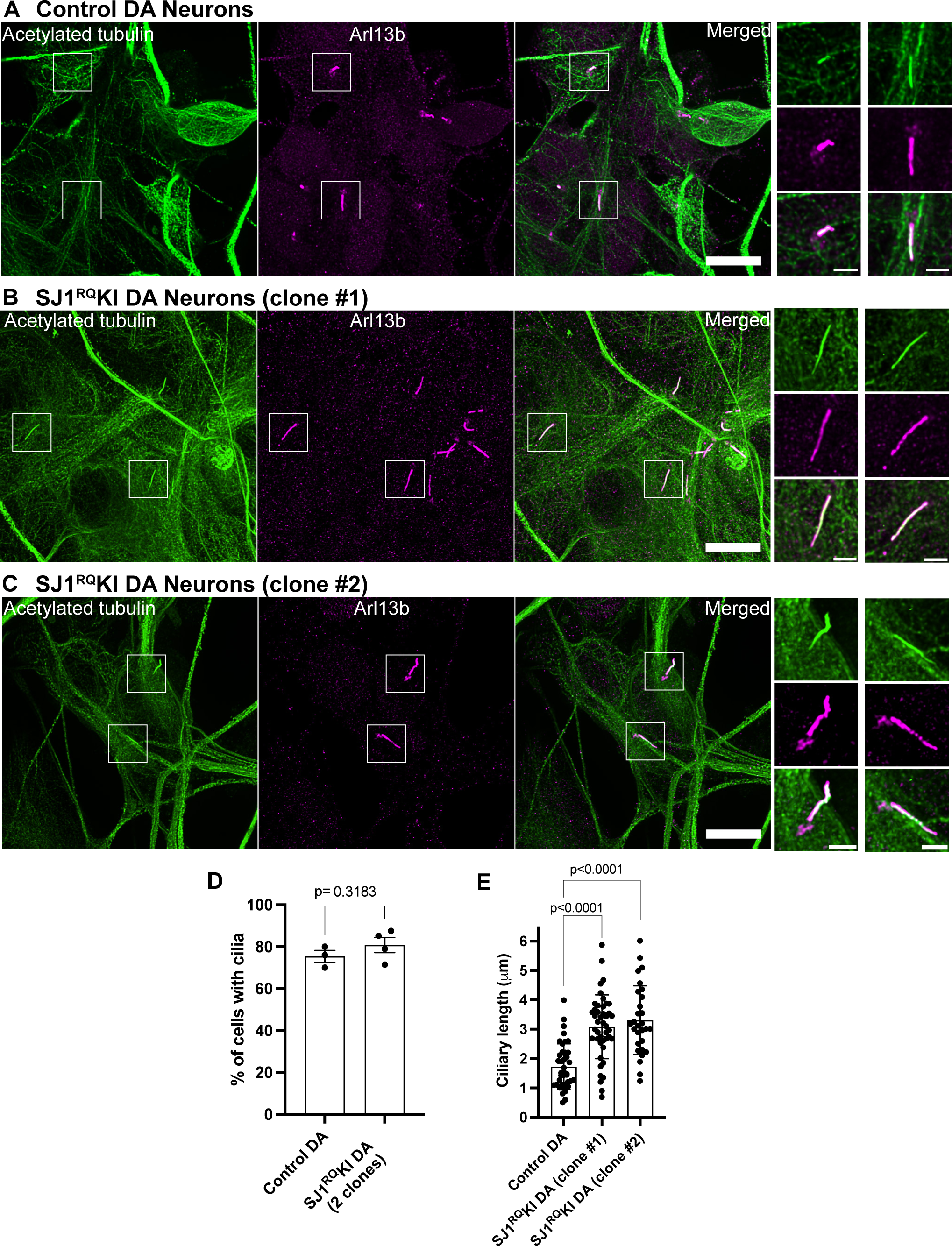
Abnormal ciliary length also in iPSC-derived SJ1^RQ^KI iPSC-derived DA neurons. (A-D) Fluorescence images of control (A) and SJ1^RQ^KI (B and C) DA neurons (day 30) derived from two KOLF2.1 iPSC clones immunolabeled with antibodies directed against acetylated α-tubulin (green) and Arl13b (magenta)(Scale bars, 10 µm). High magnifications of the boxed areas in A-C is shown on the right of each panels (Scale bars, 2 µm). (D and E) Percentage of ciliated cells (D) in control and SJ1^RQ^KI DA neurons represented as mean ± S.E.M. (from three independent experiments in which control neurons were grown in parallel with either mutant clone or both clones) (n ≥ 10 cells per experiment). (E) Ciliary length of the same control and SJ1^RQ^KI DA neurons used for panel D represented (mean ± S.D.) n ≥ 10 cells per experiment.

### Accumulation of Ca_v_1.3 in cilia of SJ1^RQ^KI DA neurons

A special property of DA neurons is an intrinsic pacemaker function, whose activity is highly dependent on the L-type Ca_v_1.2 and Ca_v_1.3 voltage-gated calcium channels (Gregory et al. 2011, Felix and Weiss 2017, Liss and Striessnig 2019, Grimaldo et al. 2022). Interestingly, these channels, which are broadly localized throughout the surface of the cell bodies and dendrites of neurons (Liss and Striessnig 2019) are also present in cilia or cilia derived structures, in several cell types, including cells of the retina and kidney (Kersten et al. 2010, Jin et al. 2014, Jin et al. 2014, Korkka et al. 2019, Sanchez et al. 2023). Prompted by this reported localization, we explored whether cilia of iPSC-derived DA neurons were labeled by anti-Ca_v_1.3 antibodies that had been validated in Ca_v_1.3 knockout cells(Shi et al. 2017). We found that in control iPSC-derived DA neurons Ca_v_1.3 immunoreactivity displayed, as previously reported (Kersten et al. 2010, Korkka et al. 2019), an accumulation at the base of cilia, whose position was marked by γ-tubulin (Figure 5A and B). Strikingly, in SJ1^RQ^KI DA neurons bright Ca_v_1.3 fluorescence intensity was observed throughout the Arl13b-positive ciliary shaft (Figure 5C-E). These findings suggest that in iPSC-derived DA neurons harboring the SJ1 PD mutation, cilia are not only abnormal in length but also in some functional properties.

**Figure 5:**
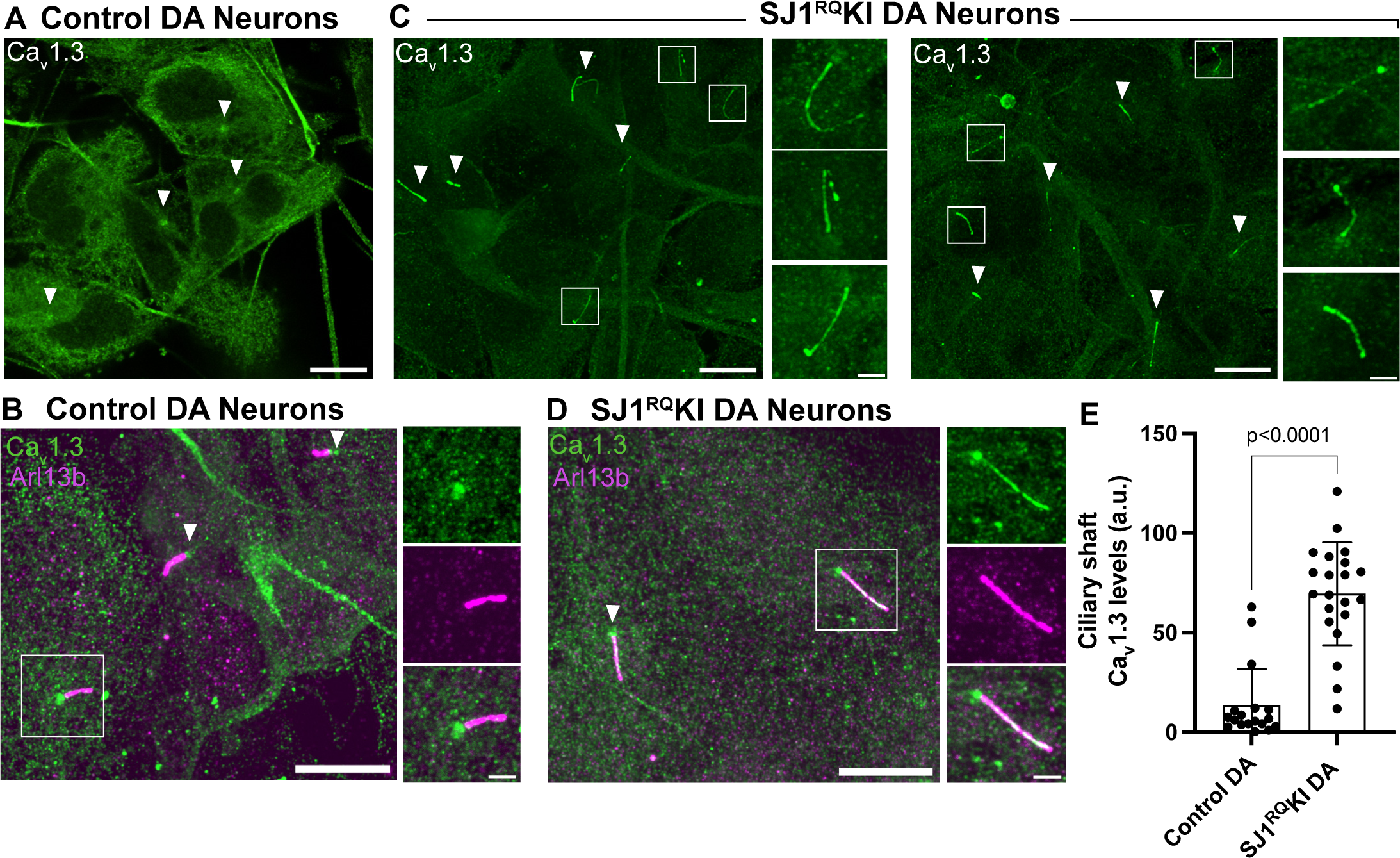
Accumulation of Cav1.3 in the ciliary shaft of SJ1^RQ^KI iPSC-derived DA neurons. (A and B) Immunofluorescence images of control iPSC-derived DA neurons demonstrating that the shaft of cilia (labeled by Arl13b; magenta) is negative for Cav1.3, which only shows some accumulation at their base (arrowheads). (C and D) Immunofluorescence images of iPSC-derived SJ1^RQ^KI DA neurons demonstrating robust labeling for Cav1.3 (green) along the shaft of Arl13b-positive (magenta) cilia. (Scale bars, 10 µm; cropped areas: 2 µm). (E) Quantification of ciliary Ca_v_1.3 immunoreactivity on the ciliary shaft of control and SJ1^RQ^KI DA neurons (mean ± S.D. pooled from three independent experiments; n ≥ 20 cells per experiment).

### Accumulation of ubiquitinated proteins in SJ1^RQ^KI DA neurons

A major mechanism underlying turnover of membrane protein in cilia is their ubiquitination, primarily via lysine 63–linked Ub (UbK63) linkage, as this process controls their exit from cilia to allow their endocytosis and targeting for degradation (Desai et al. 2020, Shinde et al. 2020, Ojeda Naharros and Nachury 2022). Thus, we investigated whether presence of ubiquitin conjugates is higher in cilia using the well-characterized FK2 and FK1 monoclonal antibodies that label ubiquitin conjugates but not free ubiquitin (Fujimuro and Yokosawa 2005)(Figure 6). While no detectable FK2 and FK1 signal was observed in the cilia of control cells, a strong signal was present in cilia of SJ1^RQ^KI DA neurons (Figure 6A-F, Supplementary Figure 3A). This result reveals a link between SJ1 function and the clearance of proteins in DA neurons, possibly reflecting back-up of endocytic traffic.

**Figure 6:**
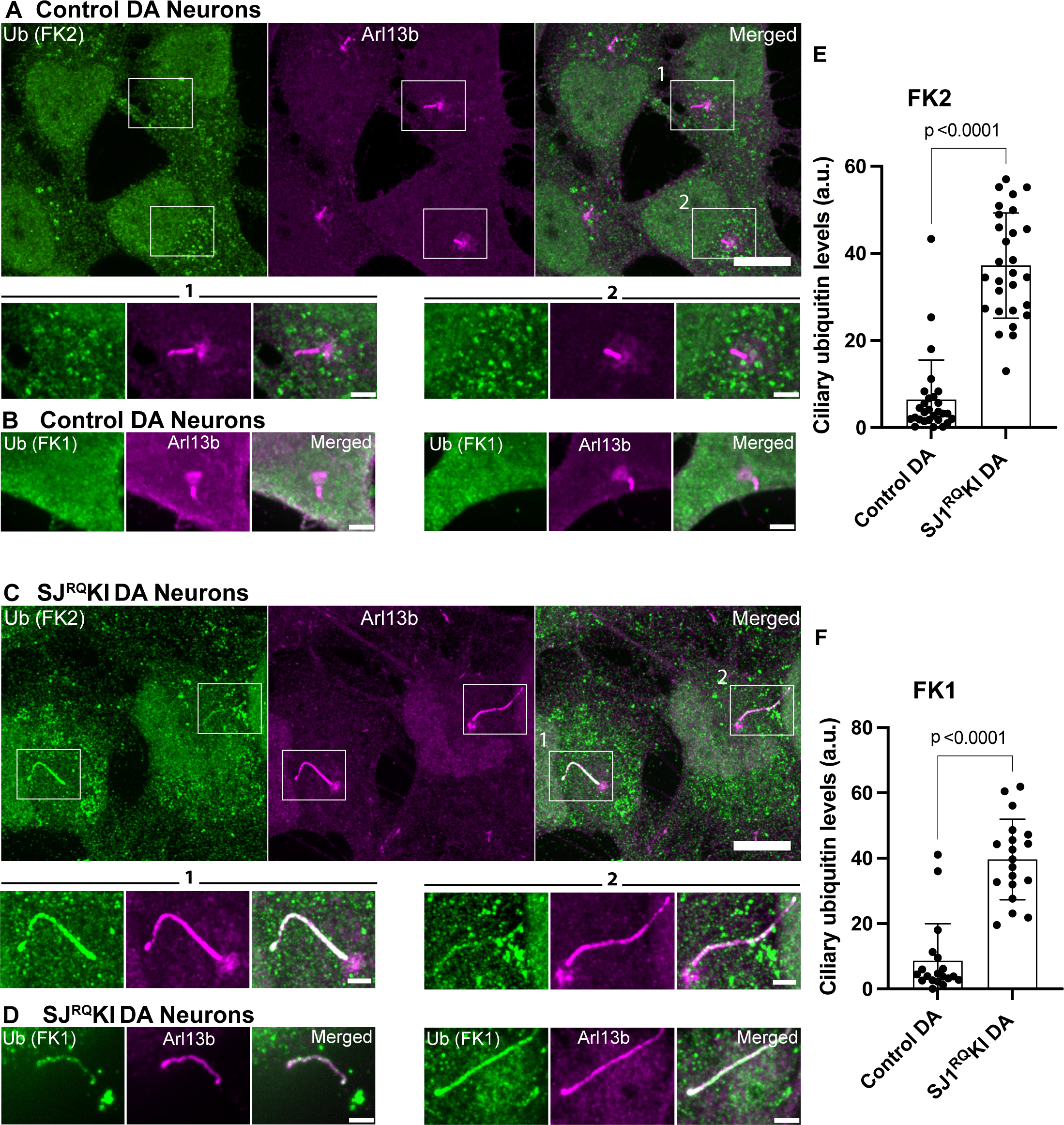
Accumulation of ubiquitin conjugates in cilia of iPSC-derived SJ1^RQ^KI DA neurons. (A-C) Fluorescence images of control (A, B) and SJ1^RQ^KI (C, D) DA neurons (day 30) immunolabeled with antibodies directed against lysine 63-linked ubiquitin chains (FK2 or FK1 antibodies, as indicated) (green) and Arl13b (magenta)(Scale bars, 10 µm). For FK2 both low and high magnifications of the boxed areas are shown (Scale bars, 2 µm) while for FK1 only high magnifications are shown. (E and F) Quantification of FK2 and FK1 immunoreactivities in the ciliary shaft of control and SJ1^RQ^KI DA neurons. Results for FK2 reflect mean ± S.D. pooled from four independent experiments (n ≥ 15 cells per experiment). Results for FK1 reflect mean ± S.D. pooled from three independent experiments (n ≥ 15 cells per experiment).

### Concentration of SJ1 at the base of primary cilia

The impact of SJ1 mutations on primary cilia could be explained by the indirect effect of an endocytic impairment throughout the neuronal surface, or to the loss of a specific function in proximity of cilia. To gain insight into this question, we assessed the localization of SJ1 by immunofluorescence in iPSCs before and after differentiation into DA neurons. We found that one or two closely apposed bright spots of SJ1 immunoreactivity colocalized with γ-tubulin, a marker of centrioles, were present in undifferentiated iPSCs and control DA neurons (Figure 7A and B). This staining at the base of cilia, was lost in SJ1 KO iPSCs and SJ1 KO DA neurons (Figure 7C and D). The localization of SJ1 at centrioles supports a role of SJ1 in cilia as it could serve as a mechanism to generate a focal high concentration of the protein in their proximity.

**Figure 7:**
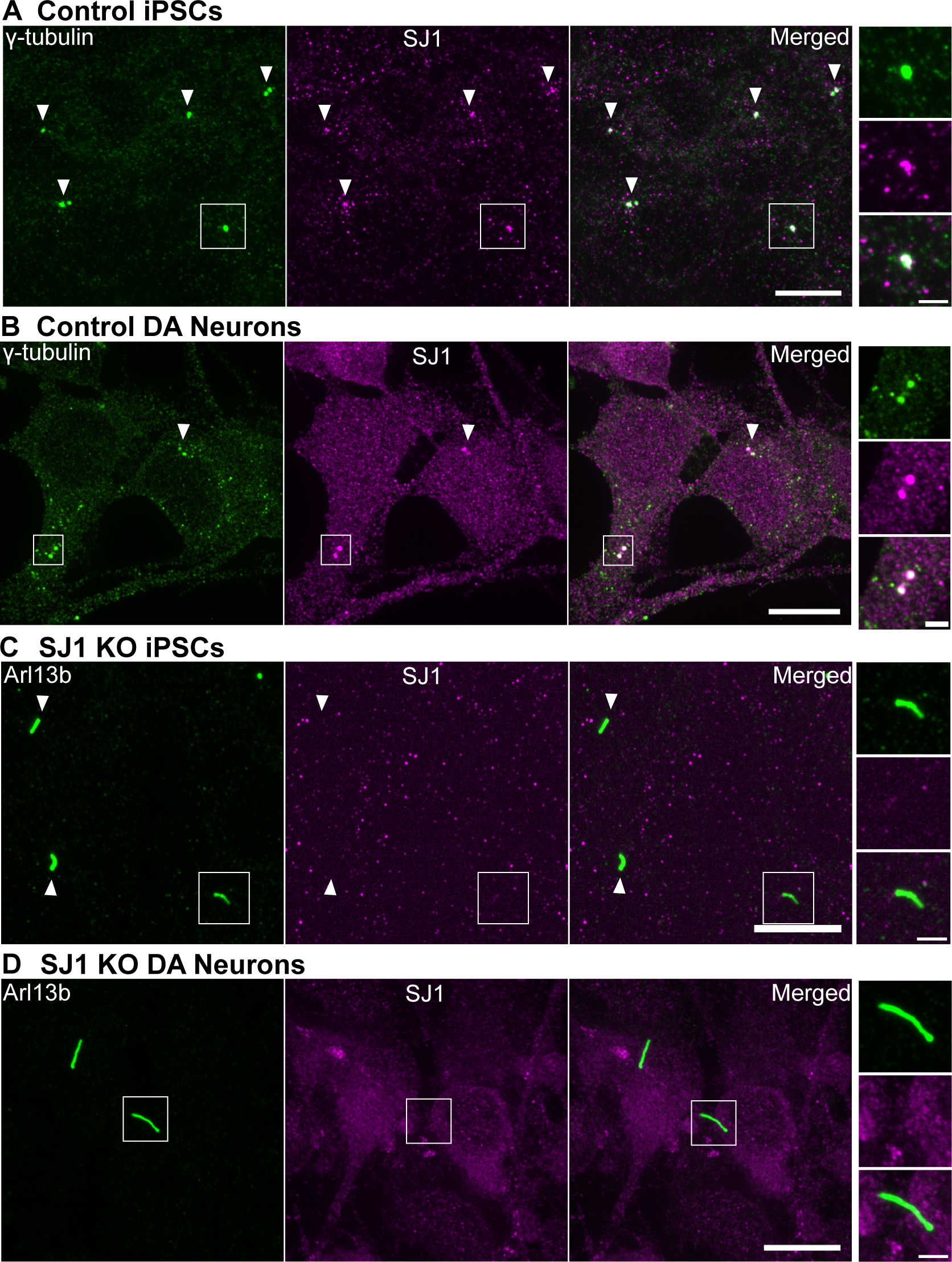
Presence of a pool of SJ1 at the ciliary base of iPSCs and iPSC-derived DA neurons. (A and B) Fluorescence images of control (A) iPSCs and (B) iPSC-derived DA neurons immunolabeled with antibodies directed against γ-tubulin (green) and SJ1 (magenta) showing overlap of spots of SJ1 immunoreactivity in control but not in SJ1 KO cells. (C and D) Fluorescence image of SJ1 KO iPSCs and iPSC-derived SJ1 KO DA neurons (day 30) immunolabeled with antibodies against Arl13b (green) and SJ1 (magenta) showing lack of SJ1 staining at the base of cilia. High magnifications of boxed areas in (A -D) are shown at right. (Scale bars, 10 µm; cropped areas: 2 µm).

A frequently used model for the analysis of cilia is the RPE1 cell line, in which serum starvation for 48 hours robustly induces ciliogenesis (Figure 8A) (Spalluto et al. 2013, Ganga et al. 2021). Upon expression of either mCherry-SJ1-145 or GFP-tagged SJ1-170 (the short and long forms of SJ1, respectively, Figure 8B and C) in these cells, bright spots of mCherry and GFP fluorescence were observed at the base of primary cilia. Co-expression in these cells of mCherry-SJ1-145 with another phosphoinositide phosphatases, the 5-phosphatase INPP5E (GFP-INPP5E), a known component of the cilia shaft (Bielas et al. 2009, Jacoby et al. 2009) confirmed the specific and selective localization of SJ1 at the cilia base (Figure 8B).

**Figure 8:**
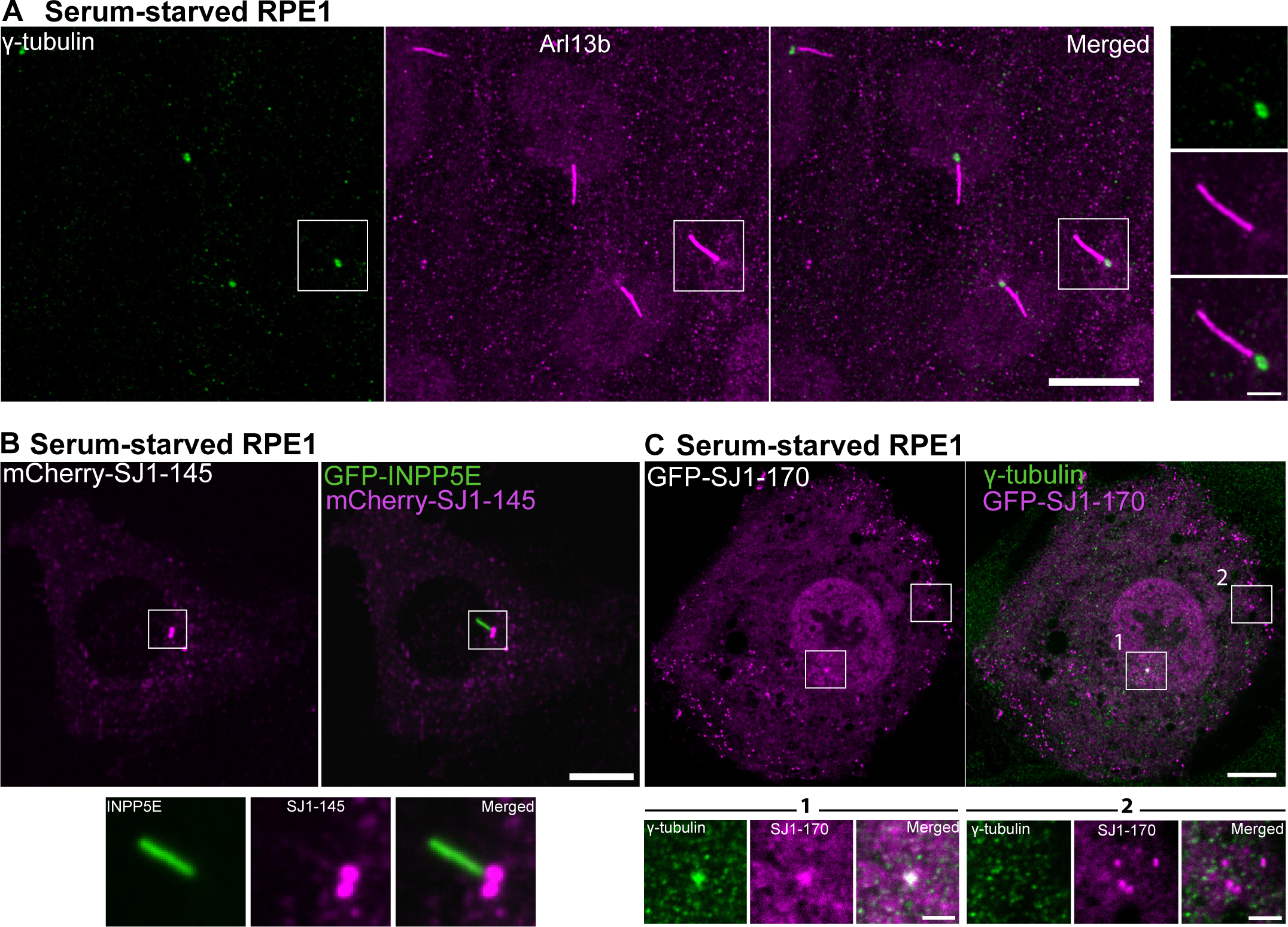
Exogenously expressed tagged-SJ1 labels the base of cilia. (A) Fluorescence image of RPE1 serum-starved for 48 hours and immunolabeled with antibodies against γ-tubulin (green) and Arl13b (magenta) show primary cilia assemblies. (B) Live fluorescence image of serum-starved RPE1 cell expressing mCherry-SJ1-145 (neuronal isoform, magenta) and GFP-INPP5E (green, a ciliary marker) showing localization of SJ1 at the ciliary base. The boxed area is shown at high magnification below the main figure. (C) Fluorescence image of serum-starved RPE1 expressing GFP-SJ1-170 (non-neuronal isoform, magenta) and immunolabeled with antibodies against γ-tubulin (green) showing overlap of the two proteins on a single perinuclear spot (boxed area 1). Boxed area 2 shows that while puncta of GFP-SJ1-170 are also observed elsewhere in the cell, these puncta do not overlap with γ-tubulin. (Scale bars, 10 µm; cropped areas: 2 µm).

## DISCUSSION

Our study shows that impairment of SJ1 function in human iPSC-derived DA neurons has an impact on the properties of their primary cilia, in addition to the well-established disrupting effect on presynaptic vesicle traffic. Both the lack of SJ1 and the selective loss of its 4-phosphatase activity due to the EOP patient mutation (SJ1^RQ^) leads to increased cilia length in these cells. Further analysis of cilia in SJ1^RQ^KI iPSC-derived DA neurons revealed abnormal protein localization in them, as exemplified by the accumulation of the Ca_v_1.3 channel and of ubiquitin chains throughout the ciliary shaft. Given the increasingly appreciated importance of primary cilia in neuronal signaling, it is plausible that a defect in ciliary function may contribute to the pathological manifestations resulting from the EOP SJ1 mutation.

Traffic of plasma membrane proteins and lipids in and out of cilia is controlled by a diffusion barrier in which PI4P (which is the predominant phosphoinositide in the ciliary shaft) and PI(4,5)P_2_ (which is the predominant phosphoinositide the ciliary pocket) play an important role (Chavez et al. 2015, Garcia-Gonzalo et al. 2015, Nguyen et al. 2022, Ojeda Naharros and Nachury 2022). Impairment of SJ1 function may disrupt the function of the diffusion barrier between the two compartments by perturbing the physiological concentration and relative ratio of PI4P and PI(4,5)P_2_. Alternatively, or in addition, SJ1 may help control membrane protein clearing from cilia indirectly via its function in the endocytic pathway (Cremona et al. 1999) as the fate of ubiquitinated proteins is to undergo endocytosis in the ciliary pocket for subsequent targeting to degradation (Nachury and Mick 2019, Shinde et al. 2020). While SJ1 appears to have a primary role in the shedding of endocytic factors after endocytosis (Nguyen et al. 2022), its loss-of-function, as shown by studies of nerve terminals, also results in a back-up of endocytic traffic with a partial stranding in the plasma membrane of proteins and membrane that needs to be internalized. As we have now shown that a pool of SJ1 is concentrated at the ciliary base, a special role of this protein in the endocytosis that occurs at the ciliary pocket is plausible.

Protein ubiquitination plays an important role in controlling protein turn-over in cilia, as a key regulatory mechanism for the exit of proteins from cilia is their ubiquitination (Anvarian et al. 2019, Nachury and Mick 2019, Shinde et al. 2020). Thus, increased cilia length and abnormal accumulation of ubiquitinated proteins in cilia may be related and due to defective protein clearance from cilia. The BBSome, a protein complex localized at cilia (Nachury et al. 2007), is implicated in this clearance (Jin et al. 2010, Chamling et al. 2014, Prasai et al. 2020). Interestingly, the BBSome components BBS7 and BBS9, as well as other proteins involved in centrosome/ciliary function, were hits in a proximity-labeling screen for SJ1 neighbors (Bartolome et al. 2022) suggesting a potential functional interplay between the BBsome and SJ1 in such clearing.

How SJ1 becomes concentrated at the base of cilia remain unclear. This localization is unlikely to be explained by its concentration on endocytic membranes in the ciliary pocket, since the localization of SJ1 closely overlaps with the localization of γ-tubulin even when the two centrioles are clearly physically separated, pointing to a concentration around the two centrioles rather than on endocytic vesicles. As the pericentriolar material is enriched in actin and actin regulatory proteins (Farina et al. 2016, Kohli et al. 2017, Kiesel et al. 2020), SJ1 may be recruited to these sites by interactions of its C-terminal proline-rich domain with actin-regulatory proteins (Rosivatz 2006). We suggest that low affinity binding to proteins that surround the centrioles may serve to create a high local concentration of SJ1, thus facilitating its action at endocytic events that takes place at these sites. We note that another inositol 5-phosphatase implicated in endocytic traffic was shown to be concentrated on centrioles at the base of cilia and impact cilia length, although with conflicting results about cilia length (Coon et al. 2012, Luo et al. 2012, Rbaibi et al. 2012), with longer cilia in Rbaibi et al. (2012).

A role in primary cilia dynamics in PD pathogenesis has been previously suggested (Nordstroma et al. 2015, Dhekne et al. 2018, Schmidt et al. 2022). In particular, at least some effects of the PD gene LRRK2 have been attributed to a role of this protein in cilia, based on studies in cell lines and mouse brain tissue (Dhekne et al. 2018, Khan et al. 2021, Sobu et al. 2021). PD mutations in LRRK2 resulted in shorter rather than longer cilia as we have shown here for SJ1 mutations. However, it remains possible that some shared aspects of ciliary function may be disrupted by both PD LRRK2 mutations and the EOP SJ1 mutation, in spite of the different effect on cilia morphology. Importantly, studies of LRRK2 and cilia have focused on striatal cells, i.e. targets of dopaminergic innervation, while here we have focused on DA neurons. An additional role of LRRK2 mutations on cilia of DA neurons cannot be excluded.

Whether and how the abnormal features of cilia of SJ1 mutant DA neurons impact their function will require further investigations. Ca^2+^ oscillations in primary cilia independent of somatic Ca^2+^ levels have been detected in several cell types and attributed to ciliary calcium channel activation, suggesting that cilia could function as an autonomous Ca^2+^ signaling hub in response to external stimuli (Delling et al. 2013, Yuan et al. 2015, Djenoune et al. 2023, Sanchez et al. 2023). In this context the striking accumulation of Ca_v_1.3 proteins in the cilia of SJ1^RQ^KI DA neurons are of special interest as it raises the possibility that Ca^2+^ signaling in these cilia may be altered, with repercussion on cell physiology.

SJ1 KO mice, which die perinatally, do not display obvious brain developmental defects at birth. Likewise, developmental defects are not observed in mice and humans with the EOP mutation. Thus, the impact of SJ1 on cilia function must be more subtle than the one of other proteins whose mutations results in major such defects, collectively referred to as ciliopathies (Reiter and Leroux 2017). Similar considerations were made for LRRK2 mutations (Dhekne et al. 2018) and OCRL mutation(Rbaibi et al. 2012).

In conclusion, our study reveals a previously unknown role of SJ1 in primary cilia of DA neurons and raises the possibility that perturbation of such role by the EOP mutation may contribute to the pathological manifestations produced by this mutation.

## ACKNOWLEDGEMENTS

This work was supported in part by grants from the National Institutes of Health (NS36251 and DA18343), and the Parkinson’s Foundation to P.D.C. This research was funded in whole or in part by Aligning Science Across Parkinson’s ASAP-000580 through the Michael J. Fox Foundation for Parkinson’s Research (MJFF). For the purpose of open access, the author has applied a CC public copyright license to all Author Accepted Manuscripts arising from this submission.

Author contributions: N.M. Rafiq and P. De Camilli conceptualized the project and wrote the manuscript. K. Fujise assisted with experiments involving non-neuronal cells and N.M. Rafiq performed everything else. P. Xu and Y. Wu provided critical research tools and M.S. Rosenfeld provided technical assistance. All authors contributed intellectually to the project.

## MATERIALS AND METHOD

### Antibodies and plasmids

mCherry-synaptojanin-145 (neuronal isoform, UniProt entry: O43426-2), GFP-synaptojanin-1-170 (non-neuronal isoform, UniProt entry: O43426) and GFP-INPP5E were previously generated in the De Camilli lab. Each construct was validated by DNA sequencing. All antibodies used in this study are listed in Table S1.

### Human iPSC culture, i^3^Neuron and DA differentiation

The following iPSC lines were obtained from the iPSC Neurodegeneration Initiative (iNDI) consortium and genome-edited by Jackson Laboratories (JAX): KOLF2.1, KOLF2.1 (with the NGN2 cassette at the AAVS locus, used for the i^3^Neurons experiments) and KOLF2.1 SJ1^RQ^KI (R219Q): clones A09 and B02. The WTC11(with the NGN2 cassette at the AAVS locus) iPSC line, kind gift of M. Ward (NIH) was used to generate SJ1 KO cells. For the maintenance of iPSCs in culture, iPSCs were cultured on Geltrex (Life Technologies) coated dishes and maintained in Essential 8 Flex media (Thermo Fisher Scientific). The Rho-kinase (ROCK) inhibitor Y-27632 (EMD Millipore, 10 μM) was added to Essential 8 Flex media on the first day of plating and replaced with fresh media without ROCK inhibitor on the following day.

For i^3^neuronal differentiation, iPSCs were differentiated into cortical-like i^3^Neurons according to a previously described protocol based on the doxycycline inducible expression of Ngn2 (Fernandopulle et al. 2018). Briefly, iPSCs were dissociated with Accutase (Thermo Fisher Scientific) and re-plated at a density between 1.5–3 × 10^5^ cells on geltrex-coated dishes in induction medium [(KnockOut DMEM/F-12 (Thermo Fisher Scientific) containing 1% N2-supplement (Thermo Fisher Scientific), 1% MEM non-essential amino acids (Thermo Fisher Scientific), 1% GlutaMAX (Thermo Fisher Scientific) and 4 μg/mL doxycycline (Sigma-Aldrich)]. After 3 days, pre-differentiated i^3^Neurons were dispersed using Accutase and plated on 0.1 mg/ml poly-L-ornithine (Sigma-Aldrich) in borate buffer and 10 μg/ml laminin (Thermo Fisher Scientific) coated 35 mm glass-bottom dishes (MatTek) or 6-well plates (Corning) for imaging and immunoblotting, respectively. These i^3^Neurons were cultured and maintained in cortical medium (induction medium supplemented with 2% B27 (Thermo Fisher Scientific), 10 ng/mL BDNF (PeproTech), 10 ng/mL NT-3 (PeproTech) and 10 μg/mL laminin). Fresh cortical media was added to the existing media every 5 days. The iPSCs and i^3^Neurons were kept at 37°C with 5% CO2 in an enclosed incubator. A detailed protocol can be found at https://www.protocols.io/view/culturing-i3neurons-basic-protocol-6-n92ld3kbng5b/v1.

For the differentiation of iPSCs to DA neurons, we used the following protocols described in Kriks et al. (2011) and Bressan et al. (2021). Briefly, iPSCs were dissociated with Accutase (Thermo Fisher Scientific) and re-plated at a density of 8 × 10_5_ cells per well (of a 6-well plate) on geltrex-coated dishes in Essential 8 Flex media with Rock inhibitor. On the next day (Day 0 of differentiation), the media was replaced with knockout serum replacement (KSR) media containing 500nM LDN193189 (STEMCELL Technologies) and 10 μM SB431542 (STEMCELL Technologies). KSR medium is comprised of Knockout DMEM/F12 medium, 15% Knockout serum replacement (Thermo Fisher Scientific), 1% MEM NEAA, 1% glutaMAX, 0.1% 2-mercaptopethanol (Thermo Fisher Scientific) and 0.2% penicillin-streptomycin (Thermo Fisher Scientific). Starting the following day (Day 1) 75% of the differentiation medium was replaced with new medium each day from day 1 to day 15, then every 2 days until day 20. For days 1-4, KSR medium containing 500 nM LDN193189, 10 μM SB431542, 200 ng/ml SHH C25II (R&D Systems), 2 μM Purmorphamine (Cayman Chemical Company) and 100 ng/ml FGF-8b (PeproTech) was added daily, supplemented by the addition of 4 μM CHIR99021 on day 3 and 4. For days 5 and 6, a mixture of 75% KSR + 25% N2 medium also containing 500 nM LDN193189, 10 μM SB431542, 200 ng/ml SHH C25II (R&D Systems), 2 μM Purmorphamine (Cayman Chemical Company), 100 ng/ml FGF-8b (PeproTech) and 4 μM CHIR99021 (Tocris) was added to the cells followed by equal amounts of KSR and N2 media on days 7-8, and 25% KSR + 75% N2 media on days 9-10 also containing 500 nM LDN193189, 10 μM SB431542, 200 ng/ml SHH C25II and 4 μM CHIR99021. The N2 medium is comprised of Neurobasal Plus media (Thermo Fisher Scientific), 2% B27 supplement without vitamin A (Thermo Fisher Scientific), 1% N2 supplement, 1% glutaMAX and 0.2% penicillin-streptomycin. For days 11-20, complete NB/B27 medium was added to cells, with the addition of 4 μM CHIR99021 on days 11 and 12 only. Complete NB/B27 medium is comprised of N2 medium (without the N2 supplement) and the following components: 20 ng/ml BDNF (PeproTech), 0.2 mM ascorbic acid (Sigma-Aldrich), 20 ng/ml GDNF (PeproTech), 0.5 mM db-cAMP (Sigma-Aldrich), 1 ng/ml TGFβ3 (R&D Systems) and 10 μM DAPT (Cayman Chemical Company). After 20 days of culture, DA progenitors cells were frozen in Synth-a-freeze cryopreservation media (Thermo Fisher Scientific) and stored at -80^0^C or liquid nitrogen.

For long-term culture of DA neurons, cells were re-plated on 0.1 mg/ml poly-L-ornithine in PBS (Sigma-Aldrich) and 10 μg/ml laminin (Thermo Fisher Scientific) coated 35 mm glass-bottom dishes (MatTek) or 6-well plates (Corning) for imaging and immunoblotting, respectively. These neurons were cultured and maintained in complete NB/B27 medium followed by the addition of 0.1% anti-mitotic inhibitor (Supplement K, Brainxell) at day 25 to terminate division of non-neuronal cells. Fresh NB/B27 medium was added to the existing plates or dishes every 7 days and kept at 37°C with 5% CO2 in an enclosed incubator.

### CRISPR–Cas9 mediated generation of SJ1 KO iPSCs

A CRISPR-based homologous recombination strategy was used to generate the SJ1 KO iPSC line. Briefly, 1 × 10^5^ WTC11-NGN2 iPSCs were plated on Geltrex-coated 6-well plate and transfected the following day using the Lipofectamine Stem transfection reagent (Invitrogen) and 3 µg of px458 plasmid (Addgene plasmid #48138) containing a small guide RNA with the following sense (5ʹC CACCGTGGTTATTACGTCTTATGTG3ʹ) and antisense (5’AAACCACATAAGACGTAATAACCAC3ʹ) sequences that was designed to selectively target the Exon 5 of SJ1. Pooled (GFP-positive) cells were enriched by fluorescence activated cell sorting (FACS) 2 days later. Sorted cells were expanded and then serially diluted to yield small clonal populations, screened using PCR amplification of genomic DNA flanking the sgRNA target site followed by sequencing of the amplicons using the following forward and reverse sequencing primers: 5’TCTCGTTTTATAGCCCTATCTTCTGATCC3’, 5’AAGGCCCATAAGTAACCAAGAACAATC3’, respectively.

### Cell culture and transfections

hTERT-RPE1 cells were grown in DMEM/F12 (Thermo Fisher Scientific) supplemented with 10% FBS (Thermo Fisher Scientific), 1% glutaMAX and 1% penicillin-streptomycin. Cells were kept at 37°C with 5% CO2 in an enclosed incubator. Cells were transfected with the relevant plasmids using 4 μls of Lipofectamine™ 2000 Transfection Reagent (Invitrogen). 4-6 hours post-transfection the medium was changed to DMEM/F12 medium without FBS to induce ciliogenesis and examined at the microscope 48 hours later. For both i^3^Neuron and DA neuron transfections, plasmids were transfected with 4 μl of Lipofectamine™ Stem Transfection Reagent (Invitrogen) and visualized at least 48 hours later.

### Immunofluorescence, live imaging and fluorescent microscopy

Cells were seeded on glass-bottom mat-tek dishes (MATtek corporation). For immunofluorescence, cells were fixed with 4% (v/v) paraformaldehyde (Electron Microscopy Sciences) in 1x phosphate-buffered saline (PBS) for 20 mins followed by three washes in PBS. Cells were permeabilized with 0.25-0.5% (v/v) Triton X-100 in PBS for 5 mins followed by three washes in PBS. Cells were then incubated with fresh 1 mg/ml sodium borohydride (Sigma-Aldrich) in PBS for 7 mins to reduce autofluorescence, and then washed thrice in PBS. They were further blocked for 30 min in 5% bovine serum albumin (BSA, Sigma-Aldrich) in PBS and then incubated overnight at 4 °C with the primary antibodies listed in Table S1. Subsequently, cells were washed with PBS thrice the following day and incubated with Alexa Fluor-conjugated secondary antibodies (Thermo Fisher Scientific) for 1 h at room temperature, followed by three washes in PBS. DAPI (Thermo Fisher Scientific) was used for nuclear staining.

Transfections were carried out as described above. For live imaging, cells were maintained in Live Cell Imaging buffer (Life Technologies) for COS7 cells, while both i^3^Neurons and DA neurons were maintained in CM and NB/B27 media, respectively, in a caged incubator with humidified atmosphere (5% CO_2_) at 37°C. The Yokogawa spinning disk field scanning confocal system with microlensing (CSU-W1 SoRa, Nikon) controlled by NIS elements (Nikon) software was used for neuronal imaging. Excitation wavelengths between 405-640 nm, CFI SR Plan ApoIR 60XC WI objective lens and SoRa lens-switched light path at 1x, 2.8x or 4x were used. SoRa images were deconvolved using the Batch Deconvolution (Nikon) software.

### Neuronal co-culture device

Control or SJ1RQKI DA neurons (day 30) were replated on one side of the two-chamber microfluidic compartmentalization device (OMEGA^4^, eNuvio), where only axonal processes can migrate through the microfluidic channels connected to the adjacent chamber. After an additional 25 days in the co-culture device, frozen iPSC-derived medium spiny neurons (MSN) from Brainxell were plated on the other half of the device (where only the axons of DA neurons are present). The DA-MSN co-cultures were then fixed 7-10 days later for immunofluorescence.

### Immunoblotting

i^3^Neurons, DA neurons and MSNs were grown on six-well plates (3-5 × 10^5^ cells/well). After differentiation in their respective maturation media, neurons were washed with ice-cold PBS and then lysed in 1xRIPA lysis buffer (10X RIPA lysis buffer, Sigma-Aldrich) supplemented with cOmplete™ EDTA-free protease inhibitor cocktail (Roche) and PhosSTOP phosphatase inhibitor cocktail (Roche), followed by centrifugation at 13,000 × g for 6 min. The supernatant was collected and incubated at 95 °C for 5 min in SDS sample buffer containing 1% 2-mercaptoethanol (Sigma). The extracted proteins were separated by SDS-PAGE in Mini-PROTEAN TGX precast polyacrylamide gels (Bio-Rad) and transferred to nitrocellulose membranes (Bio-Rad) at 100 V for 1 h or 75 V for 2 h (for high molecular weight proteins: >150 kDa). Subsequently, the nitrocellulose membranes were blocked for 1 h with 5% non-fat milk (AmericanBIO) in TBST (tris-buffered saline [TBS] + 0.1% tween 20), then incubated overnight at 4 °C with primary antibodies and then incubated with IRDye 680RD or 800CW (LI-COR) secondary antibodies (1:8000) for 1 h at room temperature in TBST. Finally, blots were imaged using the Gel Doc imaging system (Bio-Rad) using manufacturer’s protocols.

## Statistical analysis

Quantification of ciliary ubiquitination and Ca_v_1.3 levels were carried out according to Shinde et al. (2020). Briefly, total fluorescence intensity of ubiquitin or Ca_v_1.3 levels at individual Arl13b-positive cilium were substracted from background ubiquitin or Ca_v_1.3 fluorescence measured in the adjacent area. The methods for statistical analysis and sizes of the samples (n) are specified in the results section or figure legends for all quantitative data. Student’s t test or Mann– Whitney test was used when comparing two datasets. Differences were accepted as significant for P < 0.05. Prism version 9 (GraphPad Software) was used to plot, analyze and represent the data.

## Data availability

All data generated or analyzed during this study are included in this published article (and its Supplementary Information files: Supplementary Material).

---

**Table S1.** List of antibodies/dyes used in this study.

| Target Protein | Company; Catalog number | Antibody species | Working dilution for immunofluorescence | Working dilution for immunoblotting |
| --- | --- | --- | --- | --- |
| βIII-tubulin (TUJ1) | BioLegend; 801201 | Mouse | 1:500 | 1:3000 |
| βIII-tubulin | Abcam; ab18207 | Rabbit | 1:500 | N/A |
| α-Tubulin | Sigma Aldrich; T5168 | Mouse | N/A | 1:10000 |
| Tyrosine Hydroxylase | EMD Millipore; 657012 | Rabbit | 1:300 | N/A |
| Acetylated-tubulin | Sigma-Aldrich; T6793 | Mouse | 1:500 | N/A |
| ARL13b | NeuroMab; N295B/66 75-287 | Mouse | 1:250 | N/A |
| ARL13b | Proteintech;17711-1-AP | Rabbit | 1:250 | N/A |
| γ-tubulin | Sigma-Aldrich; T6557 | Mouse | 1:500 | N/A |
| Adenylate Cyclase III | Abcam; 125093 | Rabbit | 1:200 | N/A |
| Synaptojanin-1 | Novus Biologicals; NBP1-87842 | Rabbit | 1:250 | 1:1000 |
| Amphiphysin-2 | EMD Millipore; 99D | Mouse | 1:250 | N/A |
| Clathrin light chain | EMD Millipore; AB9884 | Rabbit | 1:250 | N/A |
| Auxilin | De Camilli Lab | Rabbit | 1:250 | N/A |
| Ub (FK1) | EMD Millipore; 04-262 | Mouse | 1:500 | 1:1000 |
| Ub (FK2) | Enzo Life Sciences; BML-PW8810 | Mouse | 1:500 | 1:1000 |
| Ca<sub>v</sub>1.3 | Alomone labs;ACC-005 | Rabbit | 1:250 | N/A |
| DAPI | Thermo Fischer Scientific; D3571 | N/A | 1:1000<br>(stock: 1 mg/ml) | N/A |
| Synapsin I | Synaptic Systems; 106103 | Rabbit | 1:250 | N/A |
| Synaptophysin | Synaptic Systems; 101002 | Rabbit | 1:500 | N/A |
| Synaptophysin | Synaptic Systems; 101001 | Mouse | 1:250 | N/A |
| Dopamine transporter | EMD Millipore; MAB369 | Rat | 1:500 | 1:1000 |
| Vinculin | Sigma-Aldrich; V4505 | Mouse | N/A | 1:1000 |

**Supplementary Figure 1:**
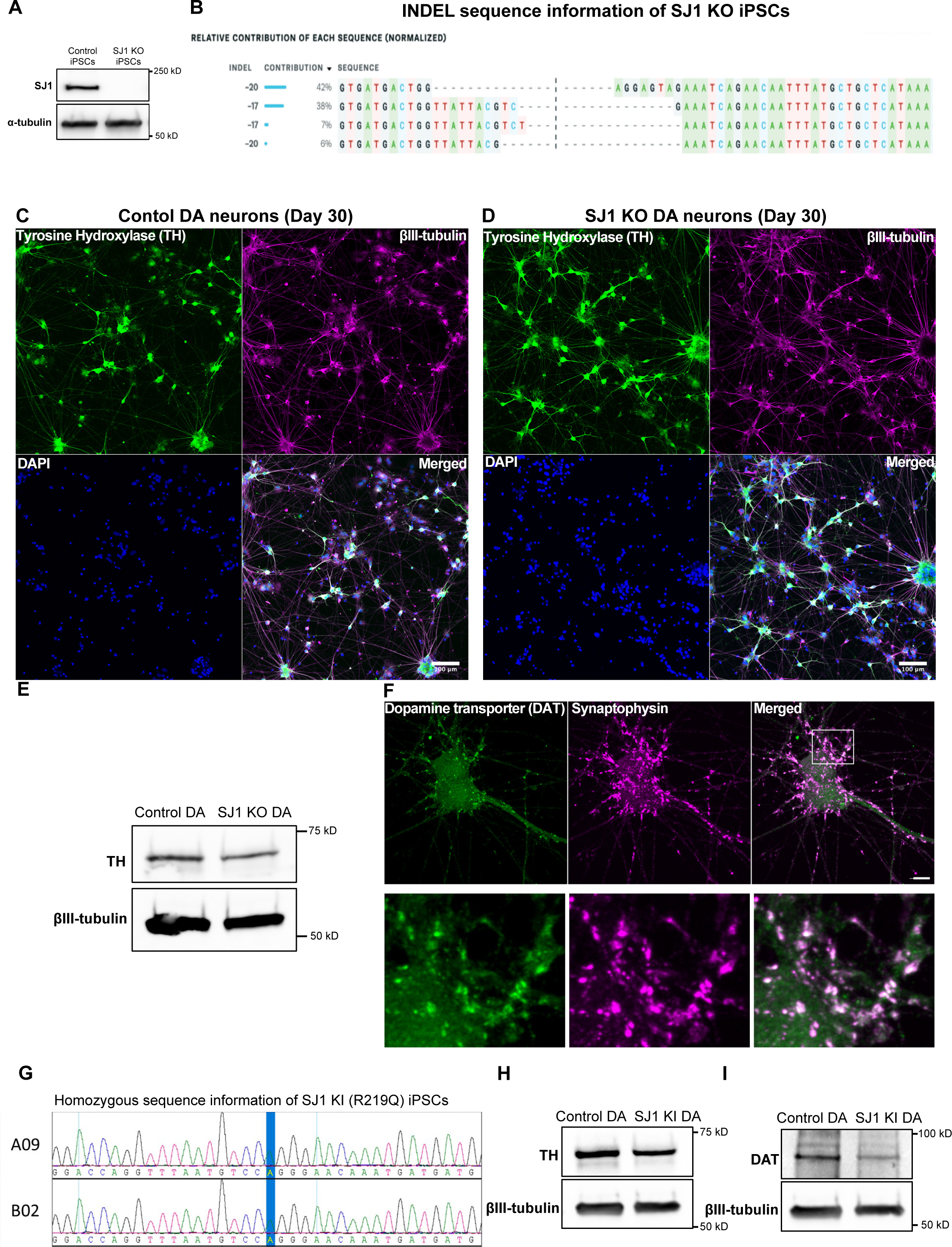
Genetic and biochemical validation of SJ1 KO and SJ1^RQ^KI iPSCs and iPSC-derived DA neurons. (A) Anti-SJ1 and anti-α-tubulin (loading control) western blot of control and edited WTC11 iPSCs. (B) Normalized relative contribution of INDEL sequences (normalized) of SJ1 KO iPSCs. (C and D) Fluorescence images of control and SJ1 KO iPSC-derived DA neurons (day 30) immunolabeled with antibodies directed against tyrosine hydroxylase (TH) (green) and βIII-tubulin (magenta). Nuclei were labeled with DAPI (blue)(Scale bars, 100 µm). (E) Anti-TH and anti-βIII-tubulin (loading control) western blot of control and SJ1 KO DA neurons (day 30). (F) Fluorescence images of control iPSC-derived DA neurons (day 30) immunolabeled with antibodies directed against dopamine transporter (DAT) (green) and synaptophysin (magenta). (G) Chromatogram sequences of two homozygous KOLF2.1 SJ1^RQ^KI iPSC clones. (H) Anti-TH and anti-α-tubulin (loading control) western blot of control and edited KOLF2.1 iPSCs. (I) Anti-DAT and anti-βIII-tubulin (loading control) western blot of control and edited KOLF2.1 iPSCs.

**Supplementary Figure 2:**
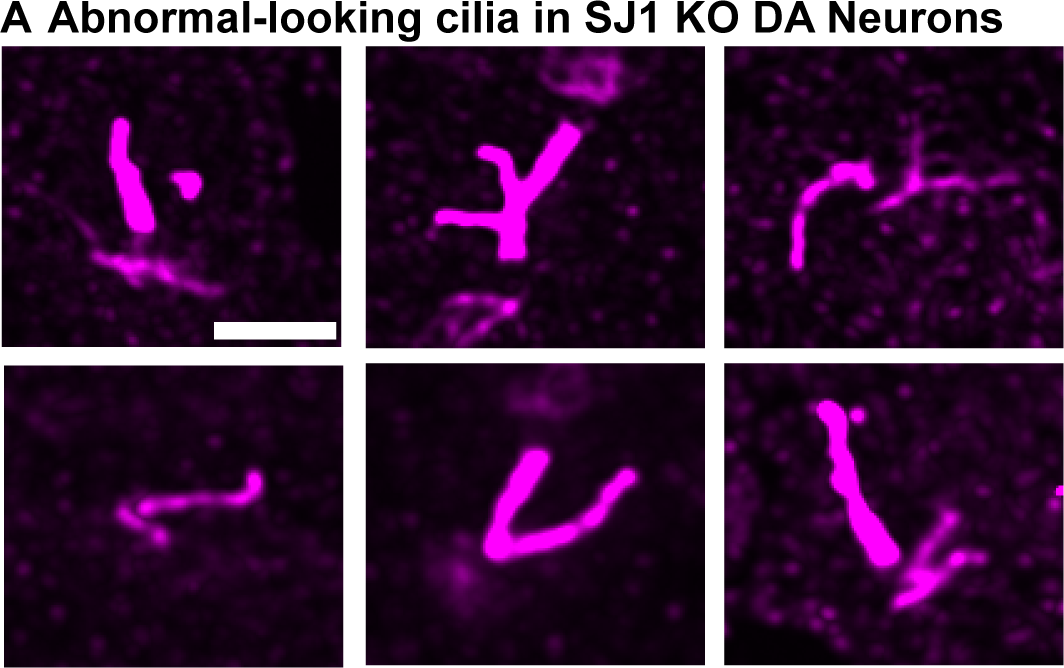
SJ1 KO DA neurons show abnormal ciliary morphology. (A) Gallery of fluorescence images of Arl13b-labeled abnormal cilia from SJ1 KO iPSC-derived DA neurons. Scale bar, 2 µm.

**Supplementary Figure 3:**
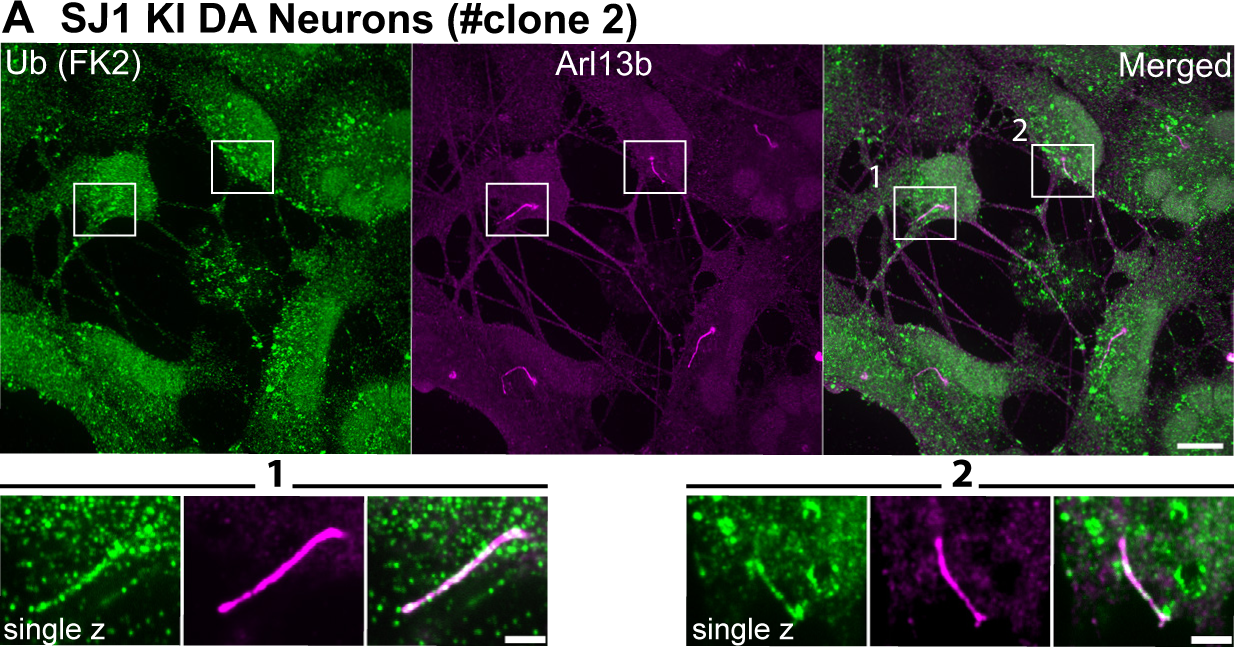
Ubiquitination in SJ1^RQ^KI DA neurons. (A) Fluorescence image of SJ1^RQ^KI (B) DA neurons clone #2 immunolabeled with FK2 antibodies directed against lysine 63-linked ubiquitin chains (green) and against Arl13b (magenta). High magnifications of the boxed areas are shown below the main panels. (Scale bars, 10 µm; cropped areas: 2 µm).

## REFERENCES

Alessi, D. R. and E. Sammler (2018). “LRRK2 kinase in Parkinson’s disease.” Science 360(6384): 36–37.

Anvarian, Z., K. Mykytyn, S. Mukhopadhyay, L. B. Pedersen and S. T. Christensen (2019). “Cellular signalling by primary cilia in development, organ function and disease.” Nat Rev Nephrol 15(4): 199–219.

Bartolome, A., J. C. Heiby, T. Dau, D. D. Fraia, I. Heinze, J. M. Kirkpatrick and A. Ori (2022). “ProteasomeID: quantitative mapping of proteasome interactomes and substrates for in vitro and in vivo studies.” bioRxiv: 2022.2008.2009.503299.

Bielas, S. L., J. L. Silhavy, F. Brancati, M. V. Kisseleva, L. Al-Gazali, L. Sztriha, R. A. Bayoumi, M. S. Zaki, A. Abdel-Aleem, R. O. Rosti, H. Kayserili, D. Swistun, L. C. Scott, E. Bertini, E. Boltshauser, E. Fazzi, L. Travaglini, S. J. Field, S. Gayral, M. Jacoby, S. Schurmans, B. Dallapiccola, P. W. Majerus, E. M. Valente and J. G. Gleeson (2009). “Mutations in INPP5E, encoding inositol polyphosphate-5-phosphatase E, link phosphatidyl inositol signaling to the ciliopathies.” Nat Genet 41(9): 1032–1036.

Blauwendraat, C., M. A. Nalls and A. B. Singleton (2020). “The genetic architecture of Parkinson’s disease.” Lancet Neurol 19(2): 170–178.

Bressan, E., A. Dhingra, S. Donato and P. Heutink (2021). “Optimized Derivation of Midbrain Dopaminergic Neurons from iPSCs for research application.” Protocols.io.

Cao, M., Y. Wu, G. Ashrafi, A. J. McCartney, H. Wheeler, E. A. Bushong, D. Boassa, M. H. Ellisman, T. A. Ryan and P. De Camilli (2017). “Parkinson Sac Domain Mutation in Synaptojanin 1 Impairs Clathrin Uncoating at Synapses and Triggers Dystrophic Changes in Dopaminergic Axons.” Neuron 93(4): 882–896 e885.

Caspary, T., C. E. Larkins and K. V. Anderson (2007). “The graded response to Sonic Hedgehog depends on cilia architecture.” Dev Cell 12(5): 767–778.

Chamling, X., S. Seo, C. C. Searby, G. Kim, D. C. Slusarski and V. C. Sheffield (2014). “The centriolar satellite protein AZI1 interacts with BBS4 and regulates ciliary trafficking of the BBSome.” PLoS Genet 10(2): e1004083.

Chavez, M., S. Ena, J. Van Sande, A. de Kerchove d’Exaerde, S. Schurmans and S. N. Schiffmann (2015). “Modulation of Ciliary Phosphoinositide Content Regulates Trafficking and Sonic Hedgehog Signaling Output.” Dev Cell 34(3): 338–350.

Coon, B. G., V. Hernandez, K. Madhivanan, D. Mukherjee, C. B. Hanna, I. Barinaga-Rementeria Ramirez, M. Lowe, P. L. Beales and R. C. Aguilar (2012). “The Lowe syndrome protein OCRL1 is involved in primary cilia assembly.” Hum Mol Genet 21(8): 1835–1847.

Cremona, O., G. Di Paolo, M. R. Wenk, A. Luthi, W. T. Kim, K. Takei, L. Daniell, Y. Nemoto, S. B. Shears, R. A. Flavell, D. A. McCormick and P. De Camilli (1999). “Essential role of phosphoinositide metabolism in synaptic vesicle recycling.” Cell 99(2): 179–188.

De Camilli, P., R. Cameron and P. Greengard (1983). “Synapsin I (protein I), a nerve terminal-specific phosphoprotein. I. Its general distribution in synapses of the central and peripheral nervous system demonstrated by immunofluorescence in frozen and plastic sections.” J Cell Biol 96(5): 1337–1354.

Delling, M., P. G. DeCaen, J. F. Doerner, S. Febvay and D. E. Clapham (2013). “Primary cilia are specialized calcium signalling organelles.” Nature 504(7479): 311–314.

Demmel, L., K. Schmidt, L. Lucast, K. Havlicek, A. Zankel, T. Koestler, V. Reithofer, P. de Camilli and G. Warren (2016). “The endocytic activity of the flagellar pocket in Trypanosoma brucei is regulated by an adjacent phosphatidylinositol phosphate kinase.” J Cell Sci 129(11): 2285.

Derderian, C., G. I. Canales and J. F. Reiter (2023). “Seriously cilia: A tiny organelle illuminates evolution, disease, and intercellular communication.” Dev Cell 58(15): 1333–1349.

Desai, P. B., M. W. Stuck, B. Lv and G. J. Pazour (2020). “Ubiquitin links smoothened to intraflagellar transport to regulate Hedgehog signaling.” J Cell Biol 219(7).

Dhekne, H. S., I. Yanatori, R. C. Gomez, F. Tonelli, F. Diez, B. Schule, M. Steger, D. R. Alessi and S. R. Pfeffer (2018). “A pathway for Parkinson’s Disease LRRK2 kinase to block primary cilia and Sonic hedgehog signaling in the brain.” Elife 7.

Di Paolo, G. and P. De Camilli (2006). “Phosphoinositides in cell regulation and membrane dynamics.” Nature 443(7112): 651–657.

Djenoune, L., M. Mahamdeh, T. V. Truong, C. T. Nguyen, S. E. Fraser, M. Brueckner, J. Howard and S. Yuan (2023). “Cilia function as calcium-mediated mechanosensors that instruct left-right asymmetry.” Science 379(6627): 71–78.

Dyment, D. A., A. C. Smith, P. Humphreys, J. Schwartzentruber, C. L. Beaulieu, F. C. Consortium, D. E. Bulman, J. Majewski, J. Woulfe, J. Michaud and K. M. Boycott (2015). “Homozygous nonsense mutation in SYNJ1 associated with intractable epilepsy and tau pathology.” Neurobiol Aging 36(2): 1222 e1221-1225.

Farina, F., J. Gaillard, C. Guerin, Y. Coute, J. Sillibourne, L. Blanchoin and M. Thery (2016). “The centrosome is an actin-organizing centre.” Nat Cell Biol 18(1): 65–75.

Felix, R. and N. Weiss (2017). “Ubiquitination and proteasome-mediated degradation of voltage-gated Ca2+ channels and potential pathophysiological implications.” Gen Physiol Biophys 36(1): 1–5.

Fernandopulle, M. S., R. Prestil, C. Grunseich, C. Wang, L. Gan and M. E. Ward (2018). “Transcription Factor-Mediated Differentiation of Human iPSCs into Neurons.” Curr Protoc Cell Biol 79(1): e51.

Fujimuro, M. and H. Yokosawa (2005). “Production of antipolyubiquitin monoclonal antibodies and their use for characterization and isolation of polyubiquitinated proteins.” Methods Enzymol 399: 75–86.

Ganga, A. K., M. C. Kennedy, M. E. Oguchi, S. Gray, K. E. Oliver, T. A. Knight, E. M. De La Cruz, Y. Homma, M. Fukuda and D. K. Breslow (2021). “Rab34 GTPase mediates ciliary membrane formation in the intracellular ciliogenesis pathway.” Curr Biol 31(13): 2895–2905 e2897.

Garcia-Gonzalo, F. R., S. C. Phua, E. C. Roberson, G. Garcia, 3rd, M. Abedin, S. Schurmans, T. Inoue and J. F. Reiter (2015). “Phosphoinositides Regulate Ciliary Protein Trafficking to Modulate Hedgehog Signaling.” Dev Cell **34**(4): 400-409.

Gregory, F. D., K. E. Bryan, T. Pangrsic, I. E. Calin-Jageman, T. Moser and A. Lee (2011). “Harmonin inhibits presynaptic Cav1.3 Ca(2)(+) channels in mouse inner hair cells.” Nat Neurosci 14(9): 1109–1111.

Grimaldo, L., A. Sandoval, P. Duran, L. Gomez Flores-Ramos and R. Felix (2022). “The ubiquitin E3 ligase Parkin regulates neuronal Ca(V)1.3 channel functional expression.” J Neurophysiol 128(6): 1555–1564.

Guo, S., L. E. Stolz, S. M. Lemrow and J. D. York (1999). “SAC1-like domains of yeast SAC1, INP52, and INP53 and of human synaptojanin encode polyphosphoinositide phosphatases.” J Biol Chem 274(19): 12990–12995.

Hardies, K., Y. Cai, C. Jardel, A. C. Jansen, M. Cao, P. May, T. Djemie, C. Hachon Le Camus, K. Keymolen, T. Deconinck, V. Bhambhani, C. Long, S. A. Sajan, K. L. Helbig, A. R. w. g. o. t. E. R. Consortium, A. Suls, R. Balling, I. Helbig, P. De Jonghe, C. Depienne, P. De Camilli and S. Weckhuysen (2016). “Loss of SYNJ1 dual phosphatase activity leads to early onset refractory seizures and progressive neurological decline.” Brain 139(Pt 9): 2420-2430.

Hulme, A. J., S. Maksour, M. St-Clair Glover, S. Miellet and M. Dottori (2022). “Making neurons, made easy: The use of Neurogenin-2 in neuronal differentiation.” Stem Cell Reports 17(1): 14–34.

Hynes, M., J. A. Porter, C. Chiang, D. Chang, M. Tessier-Lavigne, P. A. Beachy and A. Rosenthal (1995). “Induction of midbrain dopaminergic neurons by Sonic hedgehog.” Neuron 15(1): 35–44.

Jacoby, M., J. J. Cox, S. Gayral, D. J. Hampshire, M. Ayub, M. Blockmans, E. Pernot, M. V. Kisseleva, P. Compere, S. N. Schiffmann, F. Gergely, J. H. Riley, D. Perez-Morga, C. G. Woods and S. Schurmans (2009). “INPP5E mutations cause primary cilium signaling defects, ciliary instability and ciliopathies in human and mouse.” Nat Genet 41(9): 1027-1031.

Jin, H., S. R. White, T. Shida, S. Schulz, M. Aguiar, S. P. Gygi, J. F. Bazan and M. V. Nachury (2010). “The conserved Bardet-Biedl syndrome proteins assemble a coat that traffics membrane proteins to cilia.” Cell 141(7): 1208–1219.

Jin, X., A. M. Mohieldin, B. S. Muntean, J. A. Green, J. V. Shah, K. Mykytyn and S. M. Nauli (2014). “Cilioplasm is a cellular compartment for calcium signaling in response to mechanical and chemical stimuli.” Cell Mol Life Sci 71(11): 2165–2178.

Jin, X., B. S. Muntean, M. S. Aal-Aaboda, Q. Duan, J. Zhou and S. M. Nauli (2014). “L-type calcium channel modulates cystic kidney phenotype.” Biochim Biophys Acta 1842(9): 1518–1526.

Kersten, F. F., E. van Wijk, J. van Reeuwijk, B. van der Zwaag, T. Marker, T. A. Peters, N. Katsanis, U. Wolfrum, J. E. Keunen, R. Roepman and H. Kremer (2010). “Association of whirlin with Cav1.3 (alpha1D) channels in photoreceptors, defining a novel member of the usher protein network.” Invest Ophthalmol Vis Sci 51(5): 2338–2346.

Khan, S. S., Y. Sobu, H. S. Dhekne, F. Tonelli, K. Berndsen, D. R. Alessi and S. R. Pfeffer (2021). “Pathogenic LRRK2 control of primary cilia and Hedgehog signaling in neurons and astrocytes of mouse brain.” Elife 10.

Kiesel, P., G. Alvarez Viar, N. Tsoy, R. Maraspini, P. Gorilak, V. Varga, A. Honigmann and G. Pigino (2020). “The molecular structure of mammalian primary cilia revealed by cryo-electron tomography.” Nat Struct Mol Biol 27(12): 1115–1124.

Kim, T. W., J. Piao, S. Y. Koo, S. Kriks, S. Y. Chung, D. Betel, N. D. Socci, S. J. Choi, S. Zabierowski, B. N. Dubose, E. J. Hill, E. V. Mosharov, S. Irion, M. J. Tomishima, V. Tabar and L. Studer (2021). “Biphasic Activation of WNT Signaling Facilitates the Derivation of Midbrain Dopamine Neurons from hESCs for Translational Use.” Cell Stem Cell 28(2): 343–355 e345.

Klink, B. U., C. Gatsogiannis, O. Hofnagel, A. Wittinghofer and S. Raunser (2020). “Structure of the human BBSome core complex.” Elife 9.

Kohli, P., M. Hohne, C. Jungst, S. Bertsch, L. K. Ebert, A. C. Schauss, T. Benzing, M. M. Rinschen and B. Schermer (2017). “The ciliary membrane-associated proteome reveals actin-binding proteins as key components of cilia.” EMBO Rep 18(9): 1521–1535.

Korkka, I., T. Viheriala, K. Juuti-Uusitalo, H. Uusitalo-Jarvinen, H. Skottman, J. Hyttinen and S. Nymark (2019). “Functional Voltage-Gated Calcium Channels Are Present in Human Embryonic Stem Cell-Derived Retinal Pigment Epithelium.” Stem Cells Transl Med 8(2): 179–193.

Krebs, C. E., S. Karkheiran, J. C. Powell, M. Cao, V. Makarov, H. Darvish, G. Di Paolo, R. H. Walker, G. A. Shahidi, J. D. Buxbaum, P. De Camilli, Z. Yue and C. Paisan-Ruiz (2013). “The Sac1 domain of SYNJ1 identified mutated in a family with early-onset progressive Parkinsonism with generalized seizures.” Hum Mutat 34(9): 1200–1207.

Kriks, S., J. W. Shim, J. Piao, Y. M. Ganat, D. R. Wakeman, Z. Xie, L. Carrillo-Reid, G. Auyeung, C. Antonacci, A. Buch, L. Yang, M. F. Beal, D. J. Surmeier, J. H. Kordower, V. Tabar and L. Studer (2011). “Dopamine neurons derived from human ES cells efficiently engraft in animal models of Parkinson’s disease.” Nature 480(7378): 547–551.

Liss, B. and J. Striessnig (2019). “The Potential of L-Type Calcium Channels as a Drug Target for Neuroprotective Therapy in Parkinson’s Disease.” Annu Rev Pharmacol Toxicol 59: 263–289.

Luo, N., C. C. West, C. A. Murga-Zamalloa, L. Sun, R. M. Anderson, C. D. Wells, R. N. Weinreb, J. B. Travers, H. Khanna and Y. Sun (2012). “OCRL localizes to the primary cilium: a new role for cilia in Lowe syndrome.” Hum Mol Genet 21(15): 3333–3344.

McPherson, P. S., E. P. Garcia, V. I. Slepnev, C. David, X. Zhang, D. Grabs, W. S. Sossin, R. Bauerfeind, Y. Nemoto and P. De Camilli (1996). “A presynaptic inositol-5-phosphatase.” Nature 379(6563): 353–357.

Nachury, M. V., A. V. Loktev, Q. Zhang, C. J. Westlake, J. Peranen, A. Merdes, D. C. Slusarski, R. H. Scheller, J. F. Bazan, V. C. Sheffield and P. K. Jackson (2007). “A core complex of BBS proteins cooperates with the GTPase Rab8 to promote ciliary membrane biogenesis.” Cell 129(6): 1201–1213.

Nachury, M. V. and D. U. Mick (2019). “Establishing and regulating the composition of cilia for signal transduction.” Nat Rev Mol Cell Biol 20(7): 389–405.

Nemoto, Y., B. G. Kearns, M. R. Wenk, H. Chen, K. Mori, J. G. Alb, Jr., P. De Camilli and V. A. Bankaitis (2000). “Functional characterization of a mammalian Sac1 and mutants exhibiting substrate-specific defects in phosphoinositide phosphatase activity.” J Biol Chem 275(44): 34293–34305.

Ng, X. Y., Y. Wu, Y. Lin, S. M. Yaqoob, L. E. Greene, P. De Camilli and M. Cao (2023). “Mutations in Parkinsonism-linked endocytic proteins synaptojanin1 and auxilin have synergistic effects on dopaminergic axonal pathology.” NPJ Parkinsons Dis 9(1): 26.

Nguyen, T. D., M. E. Truong and J. F. Reiter (2022). “The Intimate Connection Between Lipids and Hedgehog Signaling.” Front Cell Dev Biol 10: 876815.

Nordstroma, U., G. Beauvais, A. Ghosh, B. C. Pulikkaparambil Sasidharan, M. Lundblad, J. Fuchs, R. L. Joshi, J. W. Lipton, A. Roholt, S. Medicetty, T. N. Feinstein, J. A. Steiner, M. L. Escobar Galvis, A. Prochiantz and P. Brundin (2015). “Progressive nigrostriatal terminal dysfunction and degeneration in the engrailed1 heterozygous mouse model of Parkinson’s disease.” Neurobiol Dis 73: 70–82.

Ojeda Naharros, I. and M. V. Nachury (2022). “Shedding of ciliary vesicles at a glance.” J Cell Sci 135(19).

Paisan-Ruiz, C., S. Jain, E. W. Evans, W. P. Gilks, J. Simon, M. van der Brug, A. Lopez de Munain, S. Aparicio, A. M. Gil, N. Khan, J. Johnson, J. R. Martinez, D. Nicholl, I. Marti Carrera, A. S. Pena, R. de Silva, A. Lees, J. F. Marti-Masso, J. Perez-Tur, N. W. Wood and A. B. Singleton (2004). “Cloning of the gene containing mutations that cause PARK8-linked Parkinson’s disease.” Neuron 44(4): 595–600.

Phua, S. C., S. Chiba, M. Suzuki, E. Su, E. C. Roberson, G. V. Pusapati, S. Schurmans, M. Setou, R. Rohatgi, J. F. Reiter, K. Ikegami and T. Inoue (2017). “Dynamic Remodeling of Membrane Composition Drives Cell Cycle through Primary Cilia Excision.” Cell 168(1-2): 264–279 e215.

Poewe, W., K. Seppi, C. M. Tanner, G. M. Halliday, P. Brundin, J. Volkmann, A. E. Schrag and A. E. Lang (2017). “Parkinson disease.” Nat Rev Dis Primers 3: 17013.

Prasai, A., M. Schmidt Cernohorska, K. Ruppova, V. Niederlova, M. Andelova, P. Draber, O. Stepanek and M. Huranova (2020). “The BBSome assembly is spatially controlled by BBS1 and BBS4 in human cells.” J Biol Chem 295(42): 14279–14290.

Quadri, M., M. Fang, M. Picillo, S. Olgiati, G. J. Breedveld, J. Graafland, B. Wu, F. Xu, R. Erro, M. Amboni, S. Pappata, M. Quarantelli, G. Annesi, A. Quattrone, H. F. Chien, E. R. Barbosa, N. International Parkinsonism Genetics, B. A. Oostra, P. Barone, J. Wang and V. Bonifati (2013). “Mutation in the SYNJ1 gene associated with autosomal recessive, early-onset Parkinsonism.” Hum Mutat 34(9): 1208–1215.

Ramjaun, A. R. and P. S. McPherson (1996). “Tissue-specific alternative splicing generates two synaptojanin isoforms with differential membrane binding properties.” J Biol Chem 271(40): 24856–24861.

Ramos, D. M., W. C. Skarnes, A. B. Singleton, M. R. Cookson and M. E. Ward (2021). “Tackling neurodegenerative diseases with genomic engineering: A new stem cell initiative from the NIH.” Neuron 109(7): 1080–1083.

Rbaibi, Y., S. Cui, D. Mo, M. Carattino, R. Rohatgi, L. M. Satlin, C. M. Szalinski, L. M. Swanhart, H. Folsch, N. A. Hukriede and O. A. Weisz (2012). “OCRL1 modulates cilia length in renal epithelial cells.” Traffic 13(9): 1295–1305.

Reiter, J. F. and M. R. Leroux (2017). “Genes and molecular pathways underpinning ciliopathies.” Nat Rev Mol Cell Biol 18(9): 533–547.

Rosivatz, E. (2006). “Interactions of synaptojanin.” Signal Transduction 6(2): 101–111.

Sanchez, G. M., T. C. Incedal, J. Prada, P. O’Callaghan, O. Dyachok, S. Echeverry, O. Dumral, P. M. Nguyen, B. Xie, S. Barg, J. Kreuger, T. Dandekar and O. Idevall-Hagren (2023). “The beta-cell primary cilium is an autonomous Ca2+ compartment for paracrine GABA signaling.” J Cell Biol 222(1).

Schmidt, S., M. D. Luecken, D. Trumbach, S. Hembach, K. M. Niedermeier, N. Wenck, K. Pflugler, C. Stautner, A. Bottcher, H. Lickert, C. Ramirez-Suastegui, R. Ahmad, M. J. Ziller, J. C. Fitzgerald, V. Ruf, W. D. J. van de Berg, A. J. Jonker, T. Gasser, B. Winner, J. Winkler, D. M. Vogt Weisenhorn, F. Giesert, F. J. Theis and W. Wurst (2022). “Primary cilia and SHH signaling impairments in human and mouse models of Parkinson’s disease.” Nat Commun 13(1): 4819.

Shi, L., M. L. Ko and G. Y. Ko (2017). “Retinoschisin Facilitates the Function of L-Type Voltage-Gated Calcium Channels.” Front Cell Neurosci 11: 232.

Shinde, S. R., A. R. Nager and M. V. Nachury (2020). “Ubiquitin chains earmark GPCRs for BBSome-mediated removal from cilia.” J Cell Biol 219(12).

Sipos, E., S. Komoly and P. Acs (2018). “Quantitative Comparison of Primary Cilia Marker Expression and Length in the Mouse Brain.” J Mol Neurosci 64(3): 397–409.

Sobu, Y., P. S. Wawro, H. S. Dhekne, W. M. Yeshaw and S. R. Pfeffer (2021). “Pathogenic LRRK2 regulates ciliation probability upstream of tau tubulin kinase 2 via Rab10 and RILPL1 proteins.” Proc Natl Acad Sci U S A 118(10).

Spalluto, C., D. I. Wilson and T. Hearn (2013). “Evidence for reciliation of RPE1 cells in late G1 phase, and ciliary localisation of cyclin B1.” FEBS Open Bio 3: 334–340.

Sterpka, A. and X. Chen (2018). “Neuronal and astrocytic primary cilia in the mature brain.” Pharmacol Res 137: 114–121.

Tian, R., M. A. Gachechiladze, C. H. Ludwig, M. T. Laurie, J. Y. Hong, D. Nathaniel, A. V. Prabhu, M. S. Fernandopulle, R. Patel, M. Abshari, M. E. Ward and M. Kampmann (2019). “CRISPR Interference-Based Platform for Multimodal Genetic Screens in Human iPSC-Derived Neurons.” Neuron 104(2): 239–255 e212.

Wang, C., M. E. Ward, R. Chen, K. Liu, T. E. Tracy, X. Chen, M. Xie, P. D. Sohn, C. Ludwig, A. Meyer-Franke, C. M. Karch, S. Ding and L. Gan (2017). “Scalable Production of iPSC-Derived Human Neurons to Identify Tau-Lowering Compounds by High-Content Screening.” Stem Cell Reports 9(4): 1221–1233.

Yuan, S., L. Zhao, M. Brueckner and Z. Sun (2015). “Intraciliary calcium oscillations initiate vertebrate left-right asymmetry.” Curr Biol 25(5): 556–567.

Zimprich, A., S. Biskup, P. Leitner, P. Lichtner, M. Farrer, S. Lincoln, J. Kachergus, M. Hulihan, R. J. Uitti, D. B. Calne, A. J. Stoessl, R. F. Pfeiffer, N. Patenge, I. C. Carbajal, P. Vieregge, F. Asmus, B. Muller-Myhsok, D. W. Dickson, T. Meitinger, T. M. Strom, Z. K. Wszolek and T. Gasser (2004). “Mutations in LRRK2 cause autosomal-dominant parkinsonism with pleomorphic pathology.” Neuron 44(4): 601–607.

